# A high-accuracy and high-efficiency digital volume correlation method to characterize *in-vivo* optic nerve head biomechanics from optical coherence tomography

**DOI:** 10.1101/2021.08.07.455176

**Authors:** Fuqiang Zhong, Junchao Wei, Yi Hua, Bo Wang, Juan Reynaud, Brad Fortune, Ian A. Sigal

## Abstract

*In-vivo* optic nerve head (ONH) biomechanics characterization is emerging as a promising way to study eye physiology and pathology. We propose a high-accuracy and high-efficiency digital volume correlation (DVC) method for the purpose of characterizing the *in-vivo* ONH deformation from volumes acquired by optical coherence tomography (OCT). Using a combination of synthetic tests and analysis of OCTs from monkey ONHs subjected to acute and chronically elevated intraocular pressure, we demonstrate that our proposed methodology overcomes several challenges for conventional DVC methods. First, it accounts for large ONH rigid body motion in the OCT volumes which could otherwise lead to analysis failure; second, sub-voxel-accuracy displacement can be guaranteed despite high noise and low image contrast of some OCT volumes; third, computational efficiency is greatly improved, such that the memory consumption of our method is substantially lower than with conventional methods; fourth, we introduce a parameter measuring displacements confidence. Test of image noise effects showed that the proposed DVC method had displacement errors smaller than 0.028 voxels with speckle noise and smaller than 0.037 voxels with Gaussian noise; The absolute (relative) strain errors in the three directions were lower than 0.0018 (4%) with speckle noise and than 0.0045 (8%) with Gaussian noise. Compared with conventional DVC methods, the proposed DVC method had substantially improved overall displacement and strain errors under large body motions (lower by up to 70%), with 75% lower computation times, while saving about 30% memory. The study thus demonstrates the potential of the proposed technique to investigate ONH biomechanics.

## 1. Introduction

The biomechanics of the optic nerve head (ONH) in the posterior pole of the eye play a central role in several pathologies, and are therefore important to prevent blindness [1, 2]. In glaucoma, for instance, increases in intraocular pressure (IOP) have been causally associated with higher risk of neural tissue damage and the consequent vision loss [3, 4]. The mechanisms by which higher IOP contributes to the neuropathy are not fully understood, but are known to involve IOP-induced deformations of the retinal ganglion cell axons as they pass through the ONH [5]. Improved diagnosis and treatment of glaucoma, and of other biomechanics-related ocular pathologies, therefore requires an accurate and efficient method to measure ONH biomechanics in vivo [6, 7].

Optical coherence tomography (OCT) has emerged over the last decade as the most widely used tool to image the ONH in vivo [8–17]. OCT allows acquiring three-dimensional (3D) volumes of the ONH with μm-scale resolution, and with sufficient signal penetration to visualize the lamina cribrosa region within the ONH. The lamina cribrosa is where glaucomatous neural tissue degeneration starts and is therefore of crucial interest in early diagnosis and treatment of this pathology [5,18, 19].

Also substantially advanced over the last couple of decades is the image analysis technique of digital volume correlation (DVC) [20]. DVC allows identifying and tracking corresponding points in multiple image volumes. DVC can thus be used to extract the 3D displacements and deformations between two OCT image volumes, an undeformed *reference volume*, and a *deformed volume*. The application of DVC to the analysis of OCT images of the ONH has thus emerged as a promising way to quantify the biomechanical effects caused by increases in IOP, changes in gaze, altered intracranial pressure, loss of blood pressure, and other changes in the eye biomechanical environment of potential pathologic significance [7, 21–27]. However, use of DVC for posterior pole biomechanics is hampered by the difficulty in quantifying ONH deformations from OCT volumes accurately and efficiently using existing DVC techniques originally developed other purposes. Specifically, we highlight four challenges: *First*, no two OCT scans are of precisely the same region in the exact same orientation. Thus, OCT volumes acquired at different times often exhibit both deformations and rigid body motion (*i.e*., translation and rotation) [28]. Efficient DVC requires defining an anticipated largest rigid body motion. If the rigid body motion is larger than this anticipated value, it will likely not be registered accurately. To avoid this problem it is possible to increase the size of the maximum anticipated motion, but this rapidly decreases computational efficiency. Large rotations, in particular, can also have substantial detrimental effects on the correlations, reducing the accuracy of measurements [29]. Although some motion can be reduced using motion tracking techniques, such as those in Heidelberg’s Spectralis, the movements are not entirely removed. *Second*, OCT volumes contain considerable noise compared with other imaging techniques often used for DVC. OCT scans of the ONH often have low image contrast, compounded by high speckle noise, which worsen with tissue depth further complicating analysis of deeper structures such as the lamina cribrosa, that are of great interest. Collectively, this hinders convergence of the 3D inverse-compositional Gaussian Newton (IC-GN) algorithm used in DVC [30], reducing accuracy. *Third*, conventional DVC methods are highly demanding in time and computing needs, which may make them impractical. The computational burden results primarily from the need to process a large number of points of interest (POI) within the reference volume and to do a large number of interpolations of the sub-volumes in every iteration of each POI. For example, some previous implementations of DVC for OCT have required between two and fifteen hours to track the displacements of 10,000 to 15,000 points [7, 31]. In addition, the large number of intermediary parameters needed to avoid redundant calculations increases the memory consumption. *Fourth*, interpreting DVC results is complicated as it requires simultaneously considering various aspects of the process, such as local image quality and correlation strength. In conventional DVC analyses, correlation coefficients are used implicitly to reflect the confidence of DVC tracked corresponding points. However, if the image region in question has poor contrast, a high correlation coefficient does not necessarily equate with high confidence. If points with low reliability are used to guide the DVC search path, it may compromise the robustness of the DVC computations, and the accuracy of the inferences drawn from the results [32].

In this work we present a DVC method specifically developed for application on OCT images of the ONH that addresses the four major challenges mentioned above. Our DVC method uses a combination of rigid body motion correction, sub-voxel point selection, a fast and efficient tricubic interpolation, multi-threaded parallel computations and a novel scalar measure of confidence. Herein, we describe in detail the new techniques and their implementation for the analysis of OCT volumes to quantify IOP-induced 3D deformations of the ONH. We show that the method achieves sub-voxel accuracy when registering corresponding points, in code that runs efficiently in time and computing resources. Notwithstanding its origin, the method fundamentals are not limited to this region or to OCT data, and we expect that the techniques introduced herein will prove useful for DVC in studies of other tissues.

## 2. Methods

Bellow the methods are described as per the following general order: We first describe the process of acquiring OCT volumes of the ONH in vivo at different states. The proposed DVC method is then described, in this order 1) the pre-registration technique, 2) coarse search to obtain the integer-voxel corresponding point, 3) sub-voxel registration using 3D IC-GN, 4) a fast and memory saving tricubic B-spline interpolation method, 5) confidence definition for each POI and its usage to guide the searching path and confidence weighted strain calculation, and 6) the multi-thread parallel computation technique to speed up the computation. Finally, we evaluate the DVC method on artificial volumes with various “synthetic” or preset displacements or strains and on two actual volumes captured at two controlled IOPs, and compare our method with the conventional DVC method. We should note that what we refer to as “The conventional DVC method” herein was composed of the conventional sub-voxel registration method [30], conventional tricubic B-spline interpolation method using a look-up table of control points, the conventional correlation coefficient-guided searching path, and the conventional strain calculation method, but no pre-registration [33].

### 2.1 Optic nerve head scan by an optical coherence tomography

A spectral-domain OCT device (SPECTRALIS OCT2, Heidelberg Engineering, GmbH, Heidelberg, Germany) was used to scan the ONH *in-vivo* at different states, i.e., at different longitudinal time points or under different IOP conditions. Scanning was done as described elsewhere [34]. Each OCT scan was comprised of a grid pattern of 768 x 768 A-lines, each containing 496 axial samples; the total area was 15×15 degrees and was centered on the ONH. The axial resolution in tissue is 3.87 μm and the lateral spacing between adjacent A-lines is estimated to be 5.0 μm in the monkey retina [34]. The instrument uses averaging in real time to reduce speckle noise, and these scans were obtained with a setting of N=5 sweeps averaged per B-scan. During the acquisition process, a clear, rigid gas-permeable contact lens saturated with 0.5% carboxymethylcellulose solution was placed over the apex of each cornea. Fig. 1 shows one example of the OCT acquisition of a monkey ONH. Before DVC analysis, we rescaled the OCT volumes to be isotropic, with the size of the rescaled volumes 768×770×393 voxels [24, 35].

**Fig. 1.**
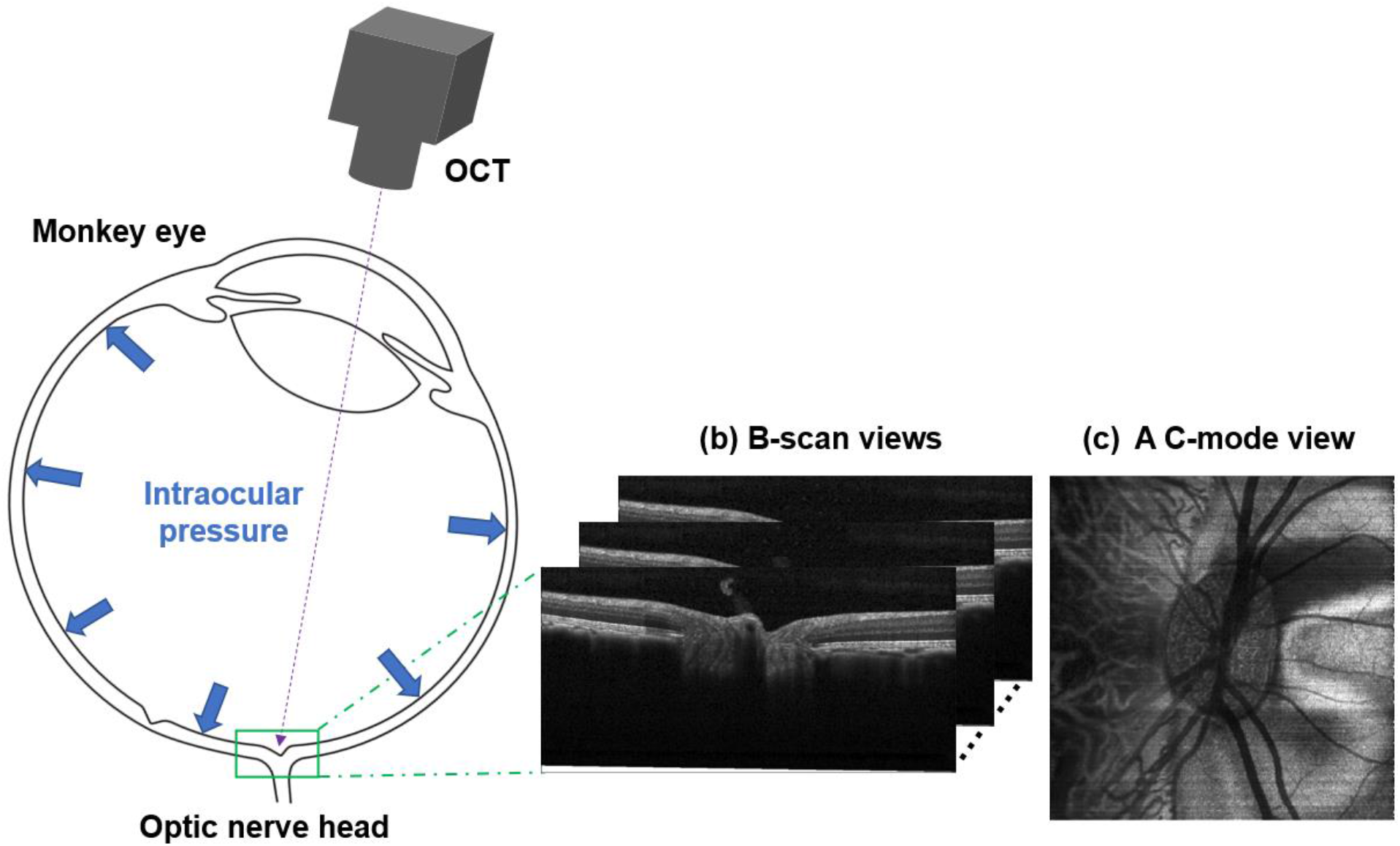
(a) Schematic longitudinal section through a monkey eye. The pressure within the globe is the intraocular pressure (IOP). Optical coherence tomography (OCT) was used to image the optic nerve head (ONH) region in the back of the eye. (b) A series of B-scans acquired from the OCT. (c) A c-mode view of the OCT volume reconstructed by these B-scans.

### 2.2 The DVC method

#### 2.2.1 A pre-registration technique to remove the rigid body motion of optic nerve head

A semi-automatic pre-registration technique combining manual operation and automatic phase correlation [36] was developed to correct the rigid body motion of the ONH between volumes, especially the rotation angles of the ONH, ensuring successful DVC correlation analysis. As shown in Fig. 2(a), interactive software with a user interface was developed to help 1) monitor the rigid body motion correction process and 2) manually adjust the position and orientation of the deformed volume to achieve registration with the reference volume. Manual operation was required if the ONH had a large rotation between the reference and deformed volumes. This approach can also help avoid converging to a spurious local optimum. To reduce the burden on the operator, the registration was done on two steps. First, the manual operation which had large error tolerance of rotation angles in the X, Y, and Z directions of about 2 degrees made the alignment step fast and easy, while already taking advantage of the operator’s experience to avoid errors. Thereafter, phase correlation was used to further correct rigid body motion automatically, and achieve an objective, precise alignment that was independent of the operator’s skill. Fig. 2(b) shows the workflow of the presented pre-registration technique. Note that the phase correlation was performed in the Fourier domain due to its high computation efficiency [37]. The translation and rotation were rectified separately in the phase correlation: correcting the translation, then the rotation. The rotation angles in the X, Y, and Z directions were also corrected independently: for example, when we correct the rotation angle *θ_z_* in the Z direction, the other two rotation angles *θ_x_* and *θ_y_* in the X and Y directions are presumed as 0. The detailed calculation process in the phase correlation is illustrated in Section 1 of Supplementary Material 1. Subsequently, a nonlinear optimization method, Nelder-Mead method, was further used to optimize (*θ_x_, θ_y_, θ_z_*) by minimizing the [*f*(*x,y,z*)~ *g*(*x*\*,*y*\*,*z*\*)]^2^, where *f* and *g* denote the reference and the deformed volumes, respectively:

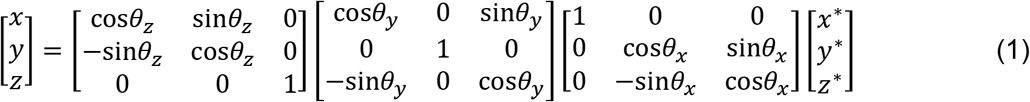

**Fig. 2.**
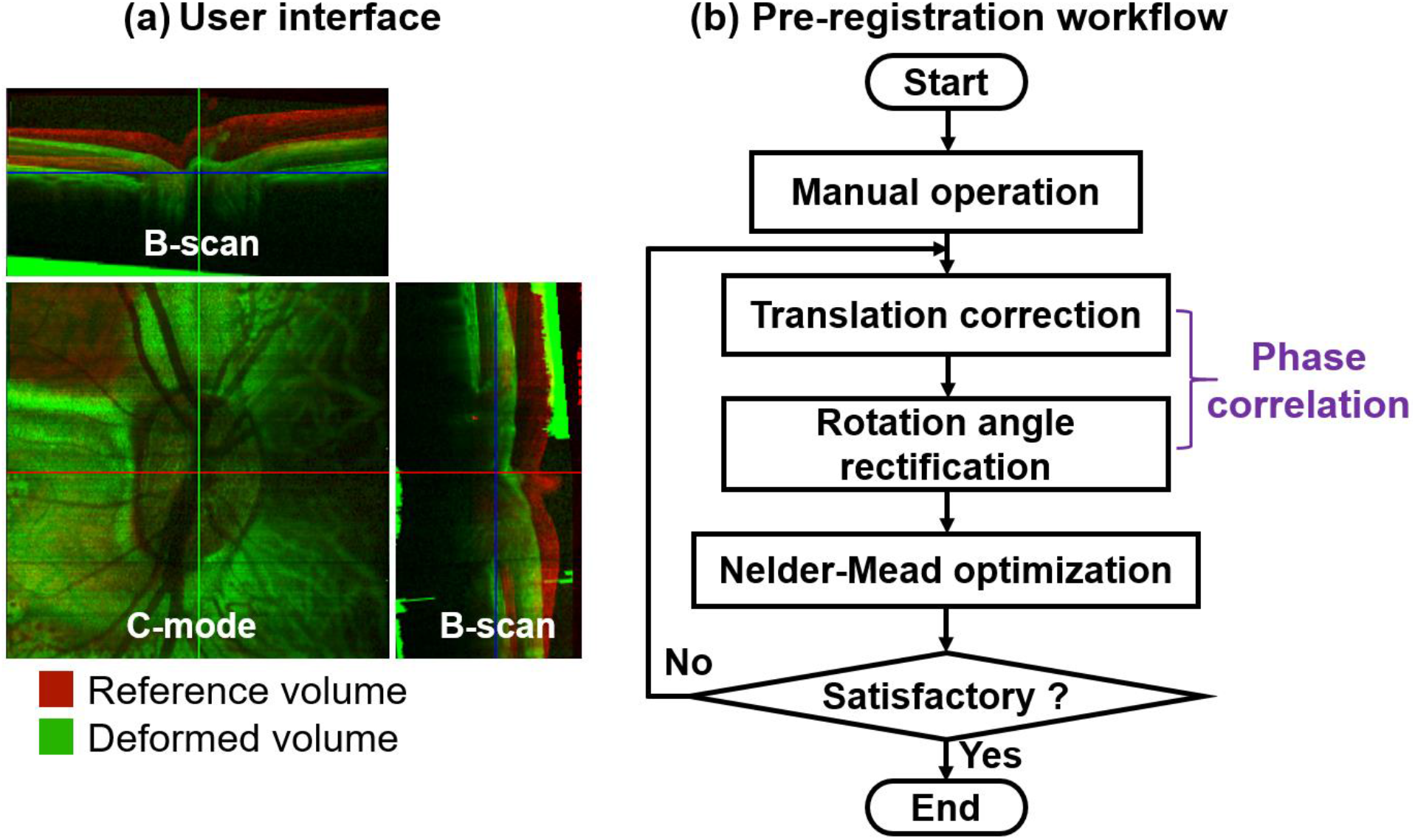
Semi-automatic pre-registration technique to correct for rigid body motion. (a) A user interface was developed to help monitor the pre-registration process and help assist with operation, illustrated with an example reference and deformed ONH volumes. The reference volume is shown in red, and the deformed volume is shown in green. The volumes are shown before registration, and thus exhibit clear displacement and rotation. (b) The workflow of the pre-registration technique. Manual operation is initially used to move and rotate the deformed volume to match with the reference volume. Phase correlation is then utilized for fine translation and rotation angle correction, followed by the Nelder-Mead optimization to optimize the rotation angle. After pre-registration we visualized the volumes. If they overall well without any easily recognized mismatches, then the registration was considered successful. If there were mismatches, we applied the process again. In our experience, we never needed more than two rounds to achieve satisfactory pre-registrations.

In practice, we can also start the next round of phase correlation to rectify the rigid body motion again until the deformed volume registers well to the reference volume, as shown in Fig. 2(b). This process is done iteratively. In our experience, one or two rounds of phase correlation were enough for most situations.

#### 2.2.2 Coarse search to obtain the integer-voxel corresponding point

As illustrated in Fig. 3, a reference subvolume with the size (2*M* + 1)^3^ voxels centered at the POI *P*_0_ was chosen in the reference volume *f*(*x, y, z*); its target subvolume was then searched pointwise in the deformed volume *g*(*x*’, *y*’, *z*’). Zero-mean normalized sum of squared difference (ZNSSD) was employed to evaluate the similarity between the reference subvolume and the searched subvolume in the deformed volume since ZNSSD is robust to the intensity linear variation, as:

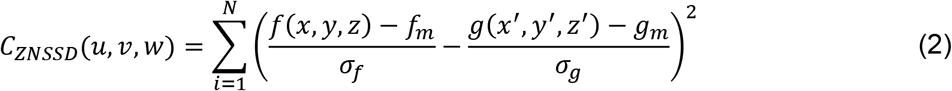

where 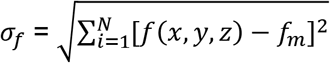 and 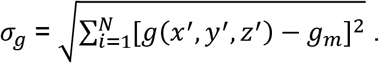 *f_m_* and *g_m_* are the average voxel intensity of the reference and deformed subvolumes, respectively. (*u,v,w*) = (*x*’,*y*’,*z*’) - (*x,y,z*). *i* is the voxel number and *N* = (2*M* + 1)^3^. The subvolume having the lowest *C_ZNSSD_* searched in the deformed volume was the target subvolume and its center 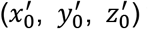 was regarded as the integer-voxel corresponding point of the POI. As a result, the initial displacement (*u*_0_, *v*_0_, *w*_0_) with integer-voxel accuracy was the difference between 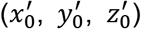 and (*x*_0_, *y*_0_, *z*_0_).

**Fig. 3.**
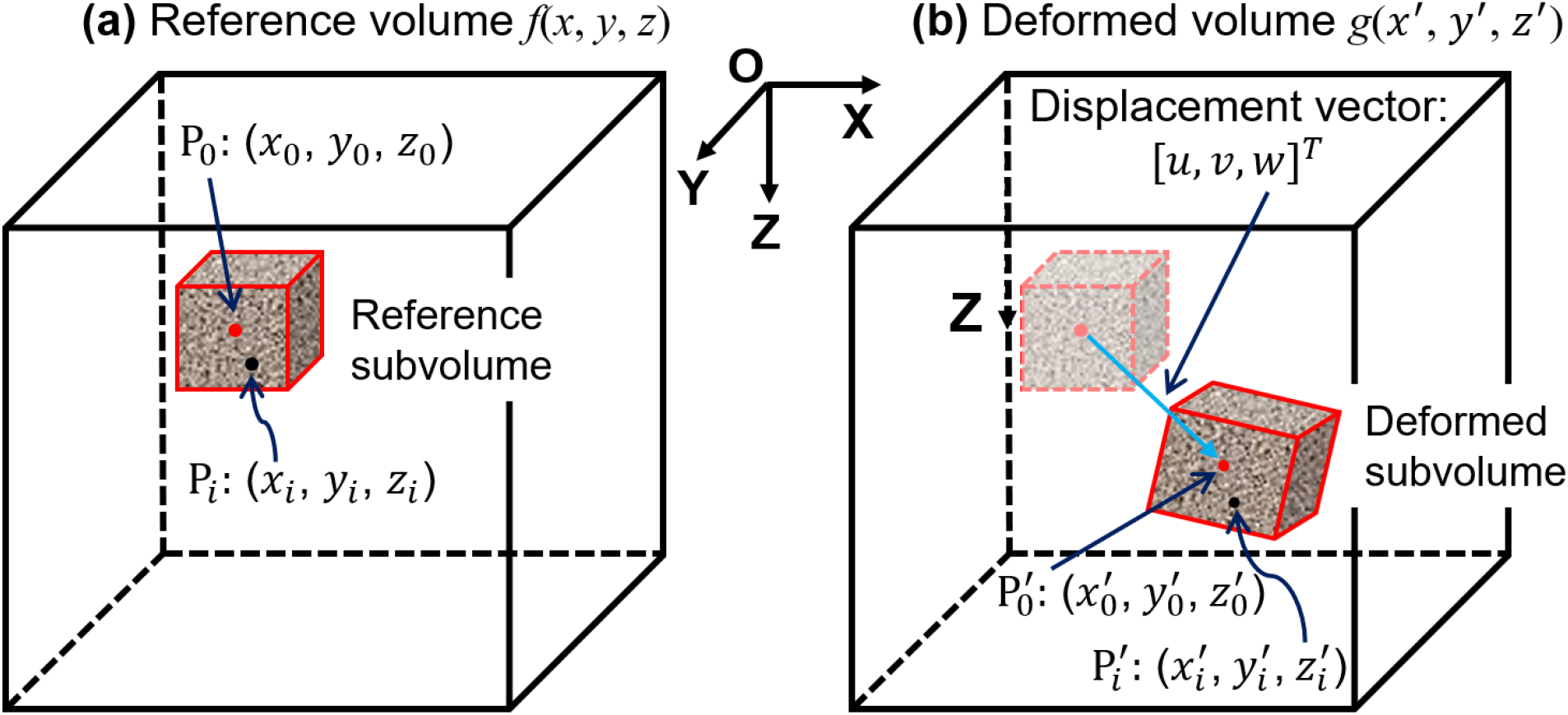
Schematic illustration of the principle of DVC which is to find the corresponding points between the reference and deformed volume by evaluating the similarity among the subvolumes. The difference between the center *P*_0_ of the reference subvolume and that center 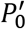 of the target deformed volume is the displacement [*u, v,w*]^*T*^. *P_i_* and 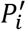 are the corresponding points in the reference and deformed subvolumes, respectively.

#### 2.2.3 Sub-voxel registration

The corresponding points from coarse search with integer-voxel accuracy were not enough to calculate the accurate strain. In this work, the popular 3D IC-GN was used to obtain the sub-voxel corresponding point in the deformed image. In the coarse search process, the shape change between the reference subvolume and the target subvolume was not considered; whereas, in the sub-voxel registration process, the first-order shape function along with the warp function **W** was employed to describe the shape change between the subvolumes by mapping a point (*x_i_, y_i_, z_i_*) around the subvolume center (*x*_0_, *y*_0_, *z*_0_) in the reference subvolume to its corresponding point 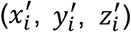 in the deformed subvolume as:

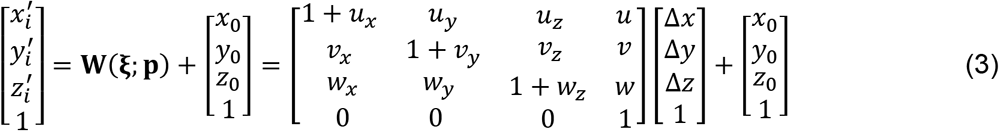

where *ξ* = [Δ*x*,Δ*y*,Δ*z*, 1]^*T*^. (Δ*x*, Δ*y*, Δ*z*) are the distance from (*x_i_, y_i_, z_i_*) to the center (*x*_0_, *y*_0_, *z*_0_). **p** represents the deformation parameters between the reference and the deformed subvolumes: **p** = {*u, v, w, u_x_, u_y_, u_z_, v_x_, v_y_, v_z_, w_x_, w_y_, w_z_*}. The first three parameters are the displacement components in the X, Y, and Z directions, respectively; while the other six are the displacement derivatives of the subvolume. Note that these six displacement derivatives in **p** are generally not as accurate as the gradients of the three displacement components to represent the deformation of the subvolume, thus, the latter was used to characterize the ONH deformation in this work. By introducing the deformation parameter **p** and the warp function **W** to consider the shape change between the reference subvolume and the target subvolume, Eq. (2) can be rewritten as

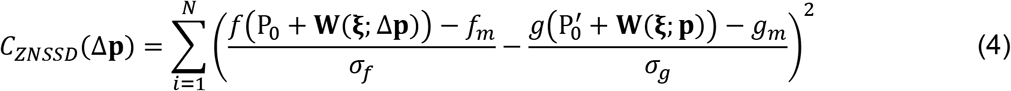

where Δ*p* is the incremental deformation parameter: Δ*p* = {Δ*u*, Δ*v*, Δ*w*, Δ*u_x_*, Δ*u_y_*, Δ*u_z_*, Δ*v_x_*, Δ*v_y_*, Δ*v_z_*, Δ*w_x_*, Δ*w_y_*, Δ*w_z_*}. *P*_0_ and 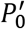 are the centers of the reference and the deformed subvolumes. The objective of this sub-voxel registration was to calculate the deformation parameter **p** by iteratively minimizing *C_ZNSSD_*. The iteration does not stop until the following convergence criterion is met:

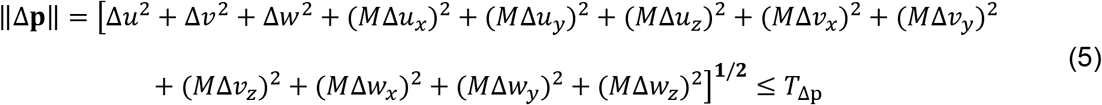

where *T*_Δ*p*_ is a threshold. In this work, when the number of iterations exceeded 20, we deemed it a failed convergence. The weak texture, low contrast, and high noise level of the OCT volume of the ONH reduce the convergence ability of the iterations. In order to ensure the accuracy, even if the iterations fail to converge, it is still required to find the sub-voxel-accuracy corresponding point. Two methods are presented in this work. The intermedia *C_ZNSSD_*, *W*(*ξ;p*) and ∥Δ*p*∥ in each iteration are recorded. Method 1 is to deduce the sub-voxel-accuracy corresponding point using the **p** having the minimal ∥Δ*p*∥. Method 2 is to do that using the **p** having the minimal *C_ZNSSD_*. The first three parameters of the corresponding **p** are the sub-voxel displacement vector. Fig. 4 shows the workflow of the 3D IC-GN iteration method.

**Fig. 4.**
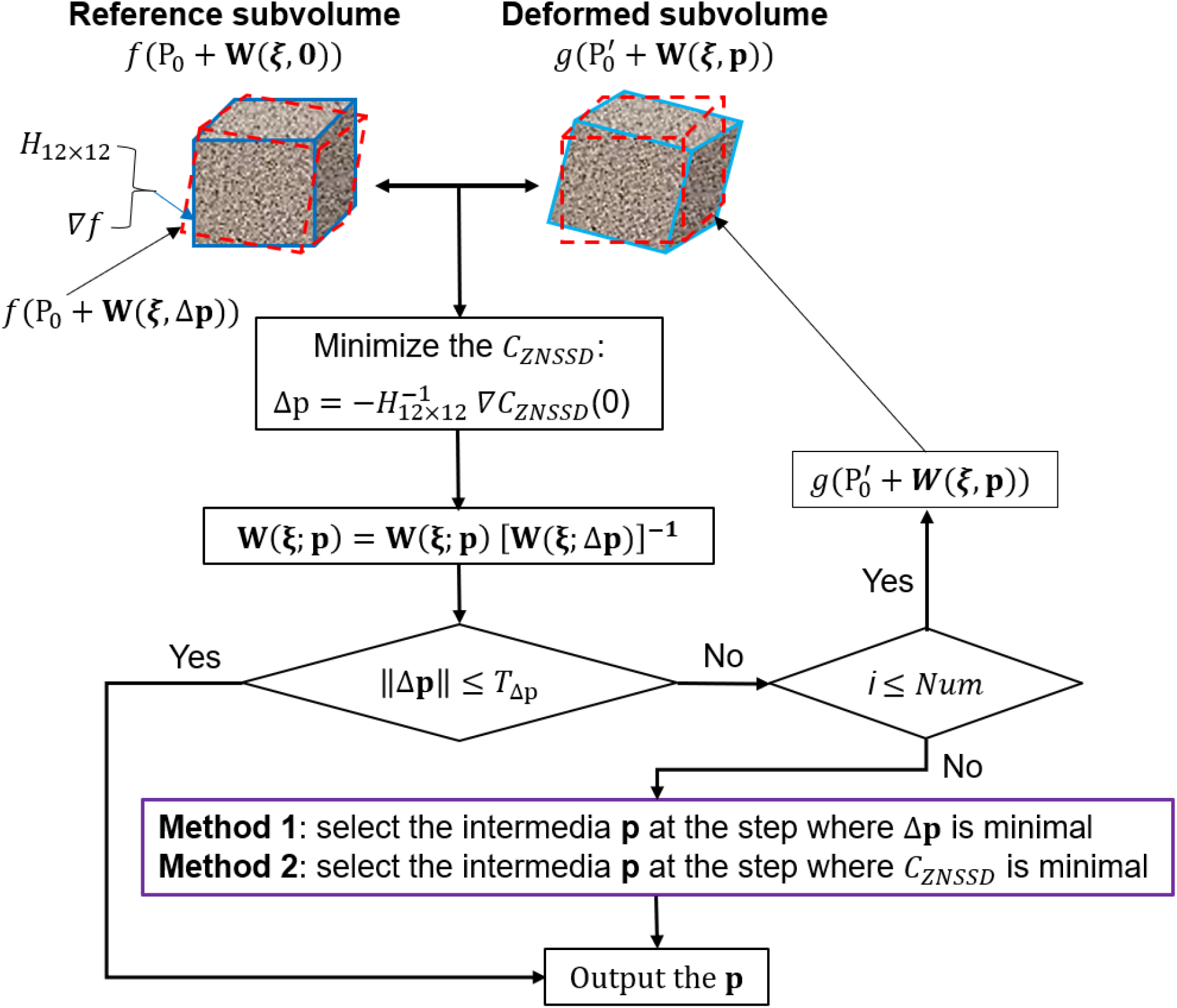
The workflow of the 3D IC-GN iteration method. Due to the weak texture and low contrast, but the large noise of the OCT volume of the ONH, the popular 3D IC-GN iteration method often fails to converge. In the conventional method, when the iteration number exceeds the limit, the corresponding point only has integer-voxel-level accuracy. We present two methods (Method 1 and Method 2) to ensure the sub-voxel accuracy when the iteration number exceeds the limit.

#### 2.2.4 A fast and memory-saving tricubic B-spline interpolation method

A large number of non-integer voxel interpolations are required to calculate the wrapped deformed subvolume in the 3D IC-GN iteration process. Due to the low accuracy of the trilinear interpolation method, in this work, the tricubic B-spline interpolation method is used to calculate the voxel intensity at the non-integer voxel position [38]. Nevertheless, the computation intensity of the conventional tricubic B-spline interpolation method is very high and it consumes a lot of memory to save the look-up table. The objective of the proposed interpolation method is to improve the computation efficiency and save the memory. As shown in Fig. 5, the presented tricubic B-spline interpolation can be divided into four bicubic B-spline interpolation and one cubic B-spline interpolation; each bicubic B-spline interpolation can further be divided into five cubic B-spline interpolation. Hence, each tricubic B-spline interpolation is composed of 21 cubic B-spline interpolations. The cubic B-spline interpolation is the basis, in which the interpolated value can be calculated by

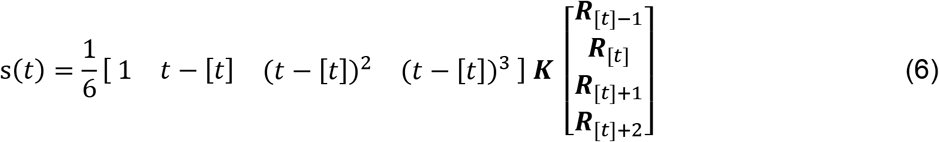

where [·] is the integer function. *t* is the position to be interpolated. ***K*** is a 4×4 interpolation kernel and it is set as [1, 4, 1, 0; −3, 0, 3, 0; 3, −6, 3, 0; −1, 3, −3, 1]. ***R*** denotes the control points determined by the discrete values ***Q*** at the regular nodes as follows:

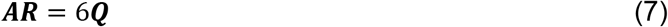

where ***A*** is the coefficient matrix. In this work, we integrate Eq. (7) into Eq. (6) by separating only four elements from ***Q*** to calculate the necessary control points ***R*** used in Eq. (6), as follows:

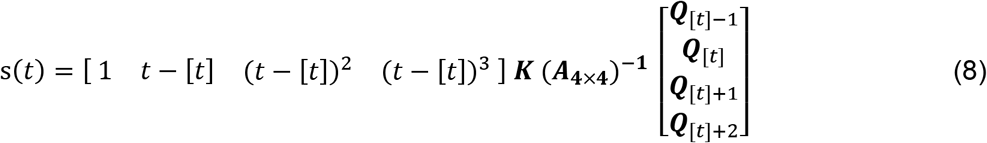

**Fig. 5.**
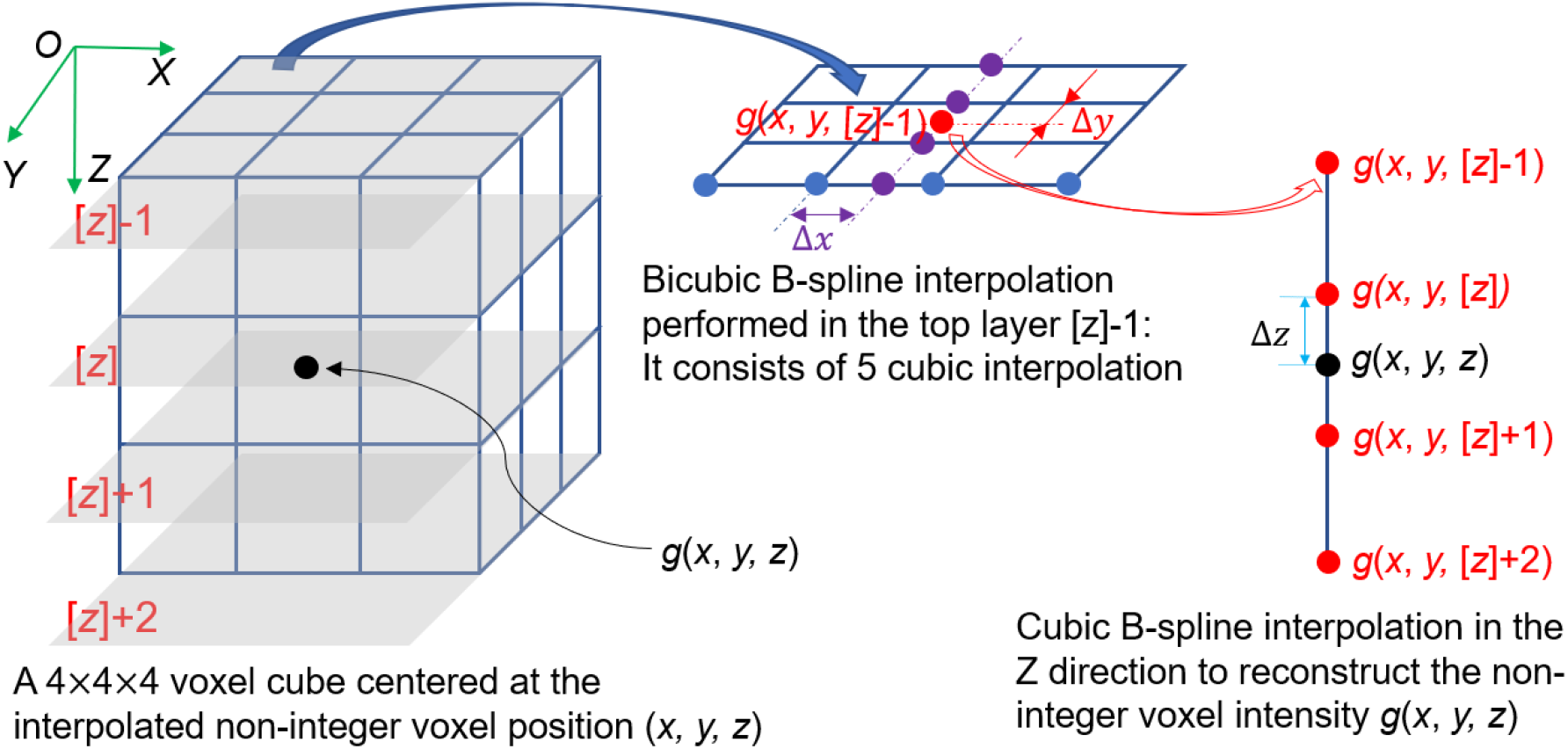
Tricubic B-spline interpolation for non-integer voxel intensity calculation. In the proposed method, it is divided into four bicubic interpolations and one more cubic interpolation. Each bicubic interpolation can also be divided into 5 cubic interpolations. Hence, one tricubic B-spline interpolation consists of 21 cubic interpolations.

If the boundary condition is set as ***R***_[*t*]-1_ = ***Q***_[*t*]-1_ and ***R***_[*t*]+1_ = ***Q***_[*t*]+1_ for each interpolation, the 4×4 coefficient matrix ***A*** becomes [6, 0, 0, 0; 1, 4, 1, 0; 0, 1, 4, 1; 0, 0, 0, 6]. In order to speed up the computation, we build up a look-up table for the results of the first three multiply factors: *LUT* = [1 *t* - [*t*] (*t* - [*t*])^2^ (*t* - [*t*])^3^] ***K*** (***A*****_4×4_**)^-**1**^. In this work, the look-up table is built when *t* - [*t*] increases from 0 to 1 at the step of Δ*t* = 0.00005. Hence, *LUT* has the size 20001 ×4 and only occupies 0.61 MB if the data is saved as *double precision* type. It is noted that *LUT* is invariant to the discrete values ***Q***, indicating no need to update the look-up table if the processed data is changed. Since one cubic B-spline in our work only includes four multiply calculation and three additive calculation, each non-integer voxel interpolation is involved with 84 multiply calculation and 63 additive calculation, indicating a relatively low computation intensity of the proposed interpolation method.

#### 2.2.5 Confidence definition for the POI and its usage to guide the searching path

In the conventional DVC method, the correlation coefficient *Corr* = 1-*C_ZNSSD_*/2 is directly used to reflect the confidence of the DVC tracked corresponding point and the larger *Corr* indicates higher confidence. The corresponding points with higher confidence are then used to guide the searching path. The position of the high reliable corresponding point can be utilized to estimate the positions of its surrounding unsearched POIs, thereby reducing the searching area greatly and enhancing the searching efficiency. However, in practice, the POI in the low contrast region even with large *Corr* sometimes has incorrect corresponding point, indicating that the large *Corr* of the POI in the low contrast region does not mean its high confidence. Using low reliable POIs to guide the searching path would negatively affect its neighboring unsearched POIs. The average voxel intensity gradient (AVIG) is a good indicator to the image contrast; the larger AVIG generally represents the higher contrast [39]. In this work, we combine both the *Corr* and AVIG to define the confidence *Conf* of the POI as follows:

If *Corr_i_* > *T_corr_* (a threshold, it set as 0.72) or *Corr_i_* < *T_corr_* & *AVIG_i_* < *Const*:

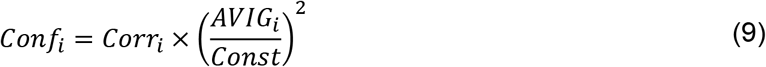

Otherwise,

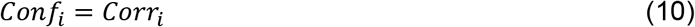

Where *Const* is a constant and set as the 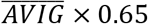 based on the experience. (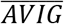 denotes the average *AVIG* of all the reference subvolumes). *i* is the subvolume number. It means that even if *Corr_i_* is large, the *Conf_i_* would still be compromised if its *AVIG_i_* is smaller than *Const*. If *Corr_i_* is smaller than the threshold *T_corr_*, it implies low reliability and *Conf_i_* would not be larger than *Corr_i_* even if its *AVIG_i_* is large. Subsequently, the POI with the *Conf* higher than a threshold *T_conf_* is then used to guide the searching path.

#### 2.2.6 Confidence-weighted strain calculation using the Savitzky-Golay filter-based method

Although rigid body motion is greatly corrected by the pre-registration technique (subsection 2.2.1), it is possible that it was not completely removed. Hence, the remaining small rigid body motion should be further removed based on the tracked corresponding points before the strain calculation [40]. The derivatives of the displacement vector in the X, Y, and Z directions are the stain components. However, the direct derivative calculation is highly sensitive to the noise, resulting in large strain error. Alternatively, in the Savitzky-Golay filter-based method (SGFM), the strains are calculated by fitting a cloud of neighboring discrete displacement vectors within a predefined cuboid (2*N*_1_+1)×(2*N*_2_+1)×(2*N*_3_+1), namely strain calculation box thereafter, using the least square method (LSM). Since the noise in the local displacement field can be greatly suppressed during the fitting process, the strains calculated in this way are much more accurate than those from the direct derivative calculation. In addition, we used the confidence *Conf* of the POI to weight every element in the displacement field. The strains 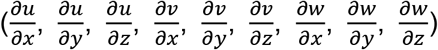 can be calculated as follows

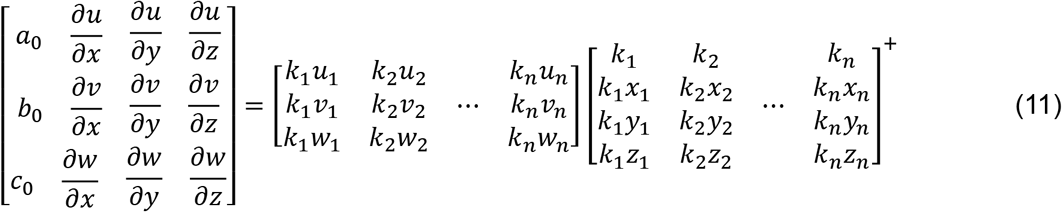

where *k_i_* = *Conf_i_*. [·]^+^ denotes the right inverse matrix. *a*_0_, *b*_0_, *c*_0_ are three parameters. (*u_i_, v_i_, w_i_*) and (*x_i_, y_i_, z_i_*) are the displacement and position of the *i*^th^ POI in the strain calculation box. The size of the strain calculation box was set as 9×9×9 in this work.

#### 2.2.7 Multi-thread parallel computation

Computation intensity of DVC is very high because the size of the subvolume for correlation is often more than 10 000 voxels and a large number of POIs have to be searched. Hence, we try to speed up the computation using the multi-thread parallel computation technique. The DVC method was programmed using C++ in Visual Studio 2019 (Microsoft Corp, Seattle, WA), except for the rigid body motion correction part, which was programmed in python. Eleven computation threads were used in our method. The workflow of the proposed DVC method using multi-thread parallel computation is shown in Section 2 of Supplementary Material 1.

### 2.3 Test methods for the proposed DVC technique

#### 2.3.1 Artificial rigid body motions to verify the pre-registration technique

An actual OCT volume of a monkey ONH was used as an example (Fig. 6). Three rigid body motions were applied to the OCT volume. Their preset translations [*T_x_, T_y_, T_z_*] were the same of [−3.2, 1.8, 9.3] voxels; while the preset rotations [*θ_x_, θ_y_, θ_z_*] were [5.3°, −5.8°, −14.4°], [−5.3°, 5.8°, 14.4°], and [−8.3°, 7.6°, −11.2°], respectively. As noted elsewhere, noise is inevitable in the actual OCT volumes. As speckle noise, a multiplicative noise, is considered the majority noise in the OCT volume [41], we added speckle noise with a standard deviation (SD) of 0.05 into each OCT volume with rigid body motion. Additionally, Gaussian noise, a common noise, was also considered in the test: Gaussian noise with the mean of 0 and the SD of 0.05 was separately added into the OCT volumes. The pre-registration technique was then used to correct the preset rigid body motions. Performance of the pre-registration technique was evaluated by the absolute difference [Δ*T_x_*, Δ*T_y_*, Δ*T_Z_*, Δ*θ_x_*, Δ*θ_y_*, Δ*θ_z_*] between the calculated rigid body motion and the preset one. Moreover, we also tested the proposed method on 7 pairs of actual OCT volumes (Supplementary Material 2).

**Fig. 6.**
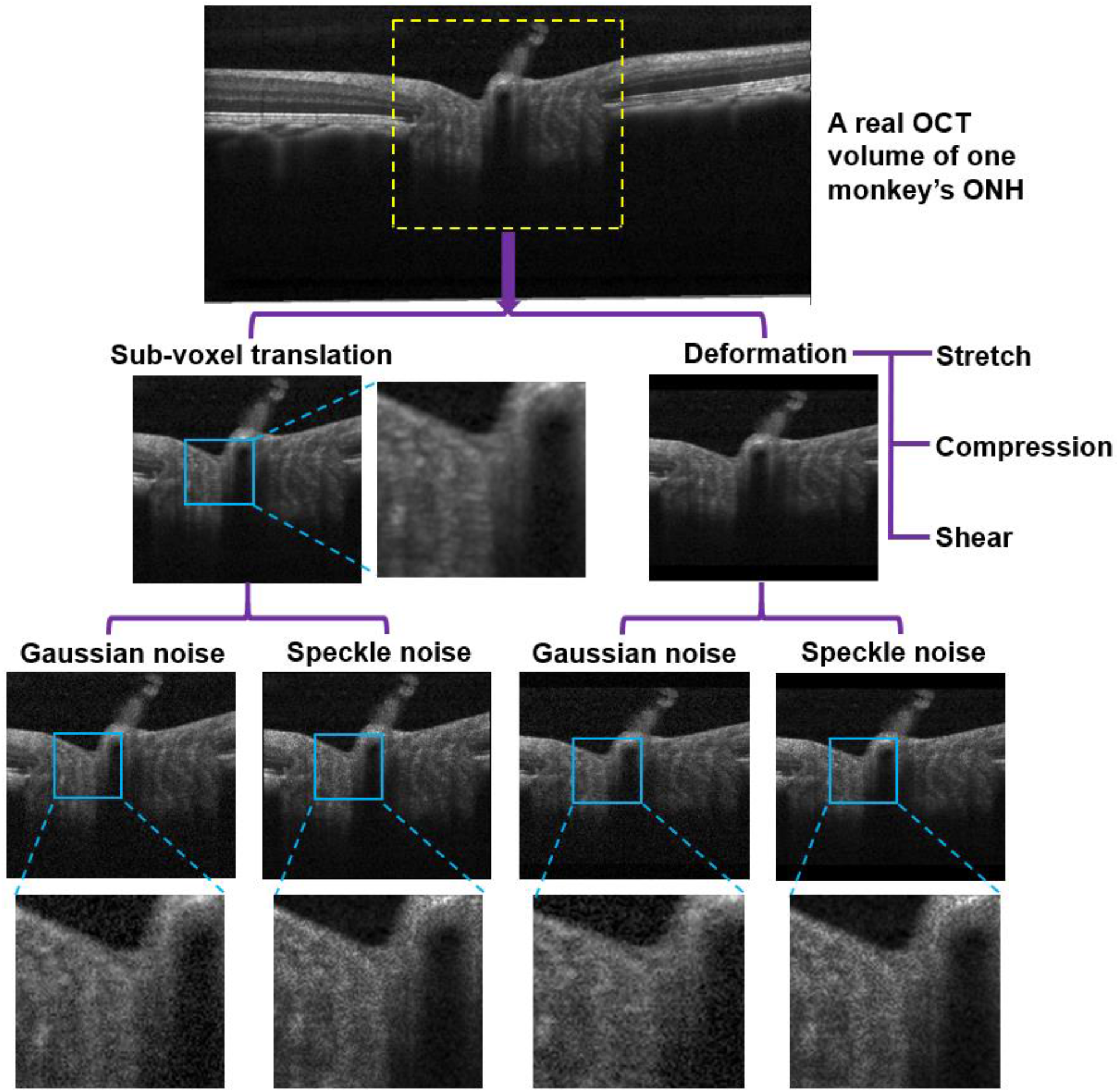
Applying rigid sub-voxel translations and various strained deformations to an actual OCT volume of a monkey’s ONH. The region enclosed by the dashed yellow frame was defined the region of interest (ROI) for this test. In addition, Gaussian noise (SD = 0.05) and/or speckle noise (SD = 0.05) are added into deformed OCT volumes. The close-ups are intended to make it easier to distinguish the noise. The rigid sub-voxel translations [*u, v, w*] in the X, Y, and Z directions are all 0.2, 0.4, 0.6, and 0.8 voxel, respectively. The strained deformations are classified into stretch, compression, and shear deformation.

#### 2.3.2 Artificial sub-voxel translations and strained deformations to test the DVC method

Various rigid sub-voxel translations and strained deformations, including stretch, compression, and shear, were applied to the actual OCT volume, as follows:

1. Rigid sub-voxel translations: *u* = *v* = *w* = 0.2, 0.4, 0.6, and 0.8 voxels, respectively.
2. Stretch: 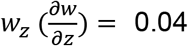, 0.07, and 0.10, respectively; while 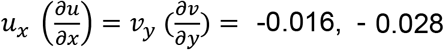 and −0.04, respectively.
3. Compression: *w_z_* = −0.04, −0.07, and −0.10, respectively; while *u_x_* = *v_y_* = 0.016, 0.028, and 0.04, respectively.
4. Shear deformation: 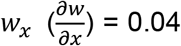, 0.07, and 0.10, respectively; while 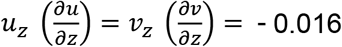, −0.028, and −0.04, respectively.
5. Shear deformation: *w_x_* = −0.04, −0.07, and −0.10, respectively; while *u_z_* = *v_z_* = 0.016, 0.028, and 0.04, respectively.

The Gaussian noise (SD = 0.05) and speckle noise (SD = 0.05) were also added to each OCT volume, respectively. The proposed DVC method was then used to measure the rigid sub-voxel translation and deformations. The accuracy and computation time of the proposed DVC method were determined.

#### 2.3.3 Comparisons between the proposed methods and the conventional methods

Several more comparisons between the proposed methods and the conventional methods were done (Supplementary Material 3):

1. the proposed sub-voxel registration methods (Method 1 and Method 2) *vs* the conventional sub-voxel registration method;
2. the proposed fast and memory saving tricubic B-spline interpolation method *vs* the existing methods to build the look-up table of the 64 interpolation coefficients or the control points;
3. the proposed confidence-guided searching method *vs* the conventional correlation-coefficient-guided searching method;
4. the proposed confidence weighted strain calculation method *vs* the conventional strain calculation method.

#### 2.3.4 Overall performance comparison

Overall performances of the proposed DVC method and conventional DVC method were compared on rigid sub-voxel translations and strained deformation measurement under rigid body motions in accuracy, efficiency, and memory consumption, as follows:

1. Rigid sub-voxel translations: *u* = *v* = *w* = 0.2, 0.4, 0.6, and 0.8 voxels, respectively.
2. Strained deformations: [*u_x_, v_y_, w_z_*] = [−0.04, −0.04, 0.1] and [0.04, 0.04, −0.1]; [*u_x_, v_y_, w_z_*] = [-0.04, −0.04, 0.1] and [0.04, 0.04, −0.1].

In either case, two kinds of rigid body motion were considered: the translations were the same of [*T_x_, T_y_, T_z_*] = [2.6, −3.4, 4.6] voxels; the rotation angles are [*θ_x_, θ_y_, θ_z_*] = [2.5°, −3.3°, 3.8°] and [−5.1°, −6.4°, 7.3°], respectively. In addition, Gaussian noise (SD = 0.05) and speckle noise (SD = 0.05) were also added to each OCT volume, respectively.

#### 2.3.5 Elevated IOP-induced deformation characterization to test the DVC method

Two OCT volumes of a monkey ONH were acquired before and after elevating the IOP from 10 mmHg to 40 mmHg and waiting for 30min. The reference volume was acquired at 10 mmHg, whereas the deformed volume was captured at 40 mmHg. The scale ratio of the two volumes: X pixel size = 4.89 μm, axial pixel size = 3.87 *μ*m, distance between scans = 4.90 *μ*m. The proposed DVC method was then applied to measure the elevated IOP-induced deformation, which was characterized by the maximal and minimal principal strains [31] as well as the maximum shear strain. The respective maximal and minimal strains are referred to as stretch and compression hereafter. The maximum shear strain is half of the absolute difference between the stretch and compression.

## 3. Results

### 3.1 Test results of the presented pre-registration technique

The absolute differences [Δ*T_x_*, Δ*T_y_*, Δ*T_z_*] between the translations measured by the pre-registration technique and the preset ones were all [0.2, 0.2, 0.3] voxels, the absolute rotation angle differences (Δ*θ_x_*, Δ*θ_y_*, Δ*θ_z_*) less than 0.4° (Table 1). Fig.7 shows an example of applying the pre-registration technique to pre-register the volume with a preset rigid body motion to its reference volume. Test of 7 pairs of OCT volumes of actual ONHs are shown in Supplementary Material 2.

**Fig. 7.**
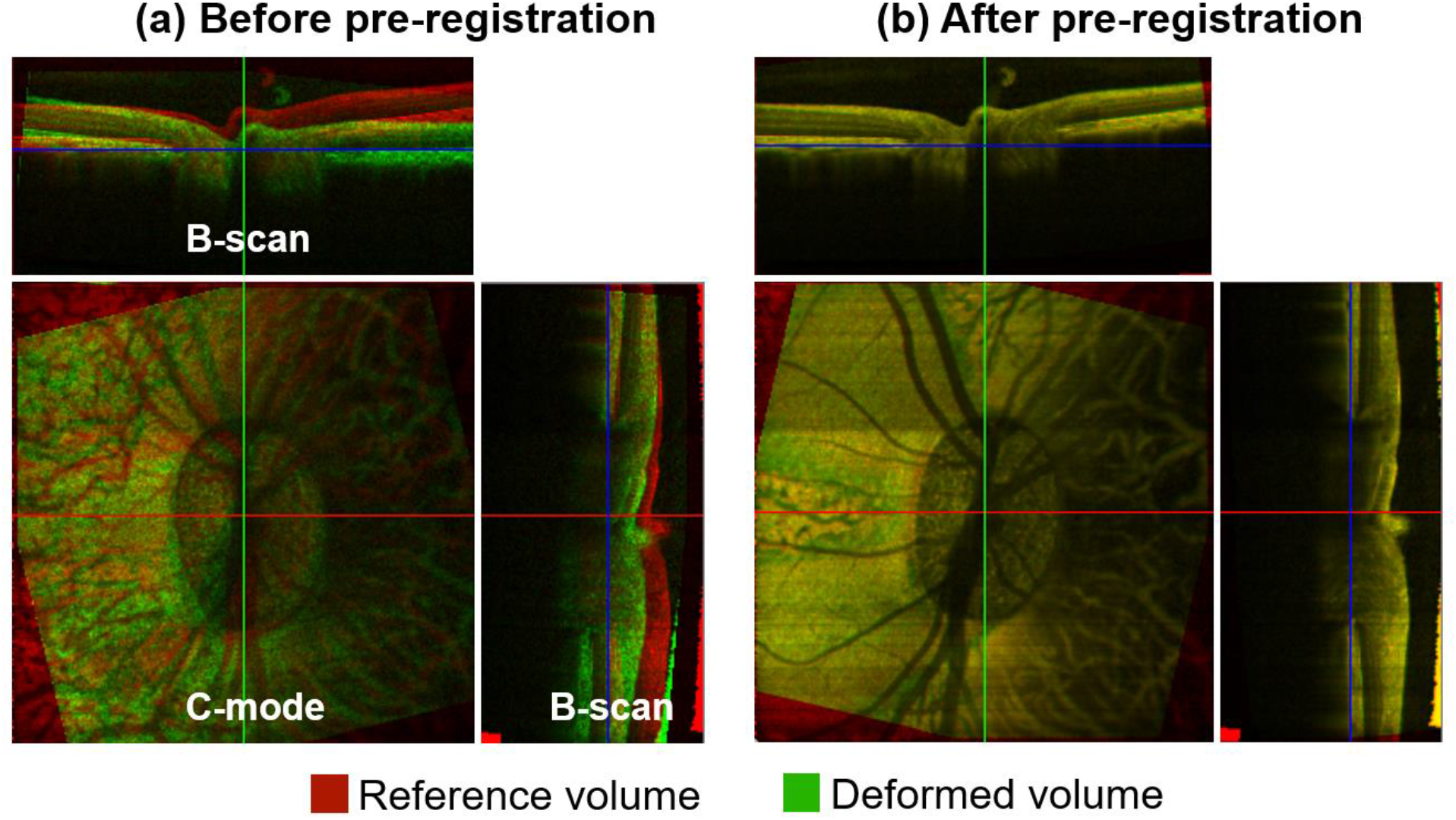
Test the pre-registration technique on rigid body motion correction. (a) The mapping before pre-registration: the red is the original OCT volume, while the green is the volume with the preset rigid body motion: [*T_x_, T_y_, T_z_, θ_x_, θ_y_, θ_z_*] = [-3.2, 1.8, 9.3, 5.3°, −5.8°, −14.4°]. The added speckle noise level is SD = 0.05. (b) The mapping after using the pre-registration technique to remove the rigid body motion. When the green voxel registers well its corresponding red voxel, it appears yellow.

**Table 1.** The absolute differences (Δ*T_x_*, Δ*T_y_*, Δ*T_z_*, Δ*θ_x_*, Δ*θ_y_*, Δ*θ_z_*) between the calculated rigid body motions and the preset ones considering Gaussian and speckle noise

|  |  |  |  |
| --- | --- | --- | --- |
| | Rigid-body motion: $(T_x, T_y, T_z) = (-3.2, 1.8, 9.3)$, $(\theta_x, \theta_y, \theta_z) =$ | | |
| | $(5.3^\circ, -5.8^\circ, -14.4^\circ)$ | $(-5.3^\circ, 5.8^\circ, 14.4^\circ)$ | $(-8.3^\circ, 7.6^\circ, -11.2^\circ)$ |
| | $(\Delta T_x, \Delta T_y, \Delta T_z, \Delta \theta_x, \Delta \theta_y, \Delta \theta_z)$ | | |
| Gaussian Noise | $(0.2, 0.2, 0.3, 0.25^\circ, 0.26^\circ, 0.36^\circ)$ | $(0.2, 0.2, 0.3, 0.16^\circ, 0.14^\circ, 0.28^\circ)$ | $(0.2, 0.2, 0.3, 0.25^\circ, 0.05^\circ, 0.33^\circ)$ |
| Speckle noise | $(0.2, 0.2, 0.3, 0.39^\circ, 0.38^\circ, 0.19^\circ)$ | $(0.2, 0.2, 0.3, 0.26^\circ, 0.18^\circ, 0.34^\circ)$ | $(0.2, 0.2, 0.3, 0.38^\circ, 0.12^\circ, 0.35^\circ)$ |

The unit of translation is voxel.

### 3.2 Test results of artificial rigid sub-voxel translations and strained deformations

Under rigid sub-voxel translations, the displacement errors in the X, Y, and Z directions (Δ*u*, Δ*v* Δ*w*) were nearly the same, under 0.037 voxel with Gaussian noise and under 0.028 voxel with speckle noise (Fig. 8(a)). Under strained deformations, average absolute strain errors in the three directions were under 0.0018 with speckle noise and under 0.0048 with Gaussian noise; average relative strain errors under 4% with Gaussian noise and under 8% under speckle noise (Fig. 8(b)). Both displacement and strain errors with Gaussian noise were larger than those with speckle noise. In addition, average absolute strain errors tended to increase with the preset deformation; whereas, the average relative strain errors had an opposite tendency. Their computation time was shown in Section 1 of Supplementary Material 3.

**Fig. 8.**
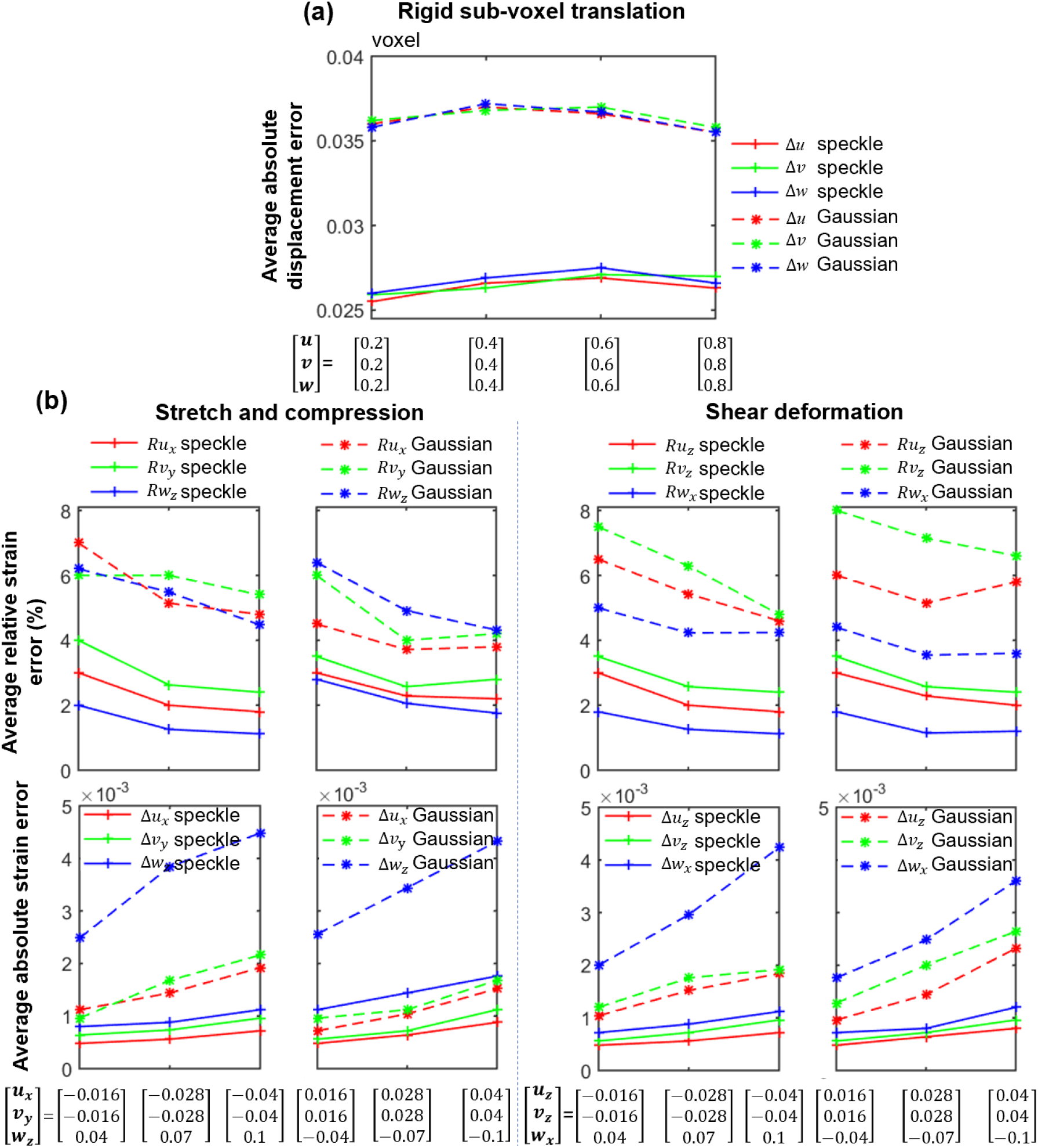
The test results of the proposed DVC method on rigid sub-voxel translations and various strained deformations in scans with Gaussian or speckle noise. (a) The average absolute displacement errors in the X, Y, and Z directions. The preset rigid sub-voxel displacements (*u, v, w*) are 0.2, 0.4, 0.6, and 0.8 voxels, respectively. (Δ*u*, Δ*v*, Δ*w*) denote the measured displacement errors in the X, Y, and Z directions, respectively. (b) The average absolute strain errors and the relative strain errors to the preset strains. Δ*u_x_* and *Ru_x_* denote the respective absolute and relative strain errors of 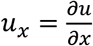 Other notations were defined in the same way. It can be observed that displacement errors in the X, Y, and Z directions are the same. Gaussian noise led to more displacement and strain errors than the same level of speckle noise. The average absolute strain errors increase with the preset strains, whereas, the relative strain errors exhibit the opposite trends.

### 3.3 Overall performance comparison results

Under rigid body motion [*T_x_, T_y_, T_z_, θ_x_, θ_y_, θ_z_*] = [2.6, −3.4, 4.6, 2.5°, −3.3°, 3.8°] (RBM1), proposed DVC had the overall displacement errors of under 0.05 voxels with speckle noise and under 0.07 voxels with Gaussian noise, and the overall strain errors of under 0.0025 with speckle noise and 0.006 with Gaussian noise; in comparison, conventional DVC method had the great larger displacement and strain errors (Fig. 9(a) and (b)). Under rigid body motion [*T_x_, T_y_, T_z_, θ_x_, θ_y_, θ_z_*] = [2.6, −3.4, 4.6, −5.1°, −6.4°, 7.3°] (RBM2), conventional DVC method failed to work, whereas, the proposed method worked well and its displacement and strain errors did not increase (Fig. 9(a) and (b)). The computation time of conventional DVC method increased from 38 minutes under rigid sub-voxel translations to more than 65 minutes under strained deformations, whereas, the computation time of proposed DVC method increased from 8 minutes under rigid sub-voxel translations to less than 18 minutes under strained deformations (Fig. 9(c)). In addition, the proposed DVC method consumed the memory of about 1.9 GB, whereas, the conventional DVC method occupied more memory of about 2.8 GB. The separate comparison results between the proposed methods and conventional methods are demonstrated in Supplementary Material 3.

**Fig. 9.**
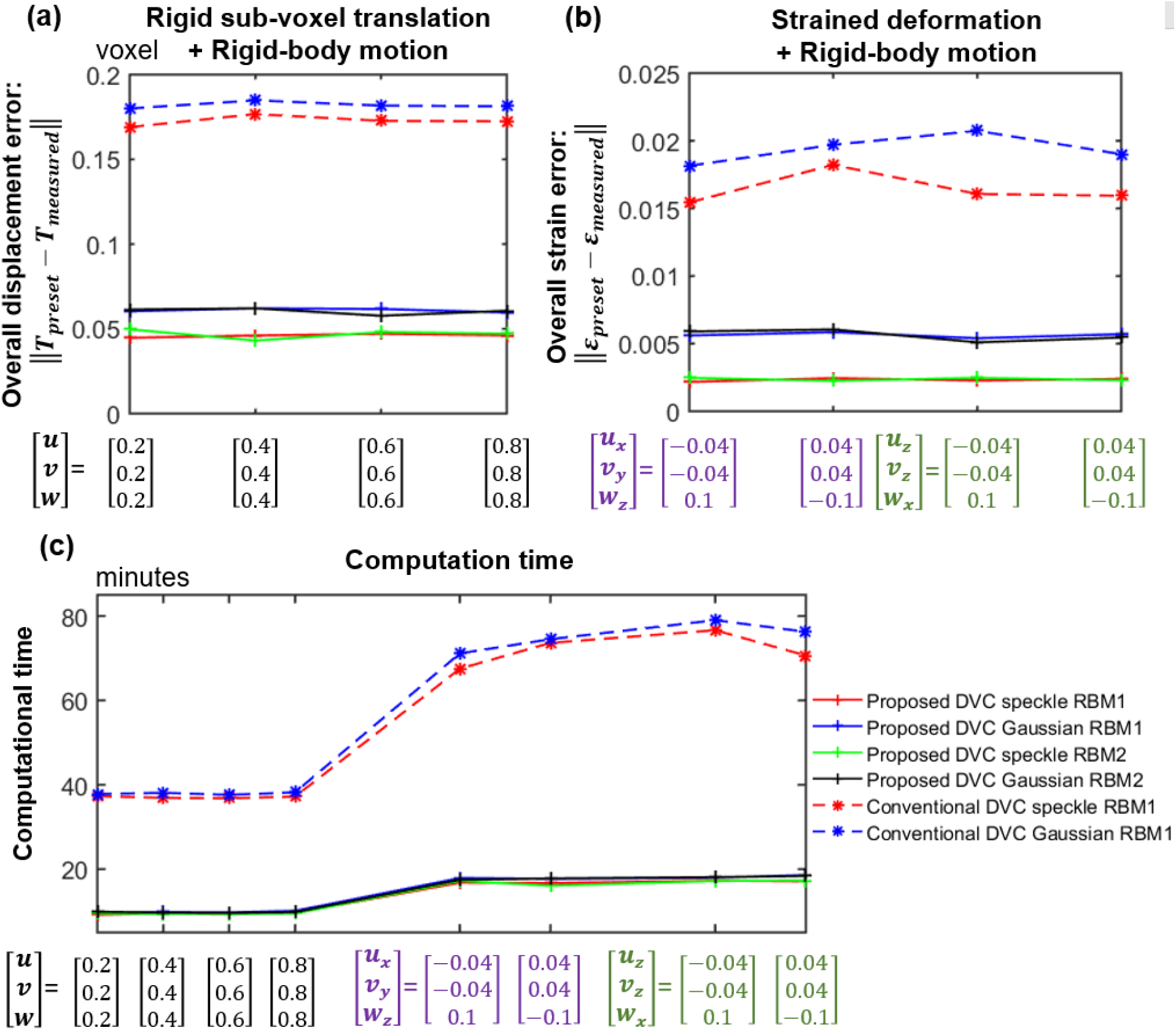
Overall performance comparison between the proposed DVC method and the conventional DVC method on displacement and strain measurement accuracy and computation time under rigid body motion. Two kinds of rigid body motion (RBM1 and RBM2) are set. RBM1: [*T_x_, T_y_, T_z_, θ_x_, θ_y_, θ_z_*] = [2.6, −3.4, 4.6, 2.5°, −3.3°, 3.8°]; RBM2: [*T_x_, T_y_, T_z_, θ_x_, θ_y_, θ_z_*] = [2.6, −3.4, 4.6, −5.1°, −6.4°, 7.3°]. (a) The overall displacement errors, *T_preset_* can be deduced from the preset rigid sub-voxel translation and rigid body motion. (b) The overall strain errors. Note that the conventional DVC method failed to work in RMB2, and thus its results are not shown. The results of the proposed DVC method under RBM1 and RBM2 were very similar. It can be seen that, in case of rigid body motion, the proposed DVC method is clearly superior to conventional DVC method in overall displacement error, overall strain error, and computation time.

### 3.4 Results of the elevated IOP-induced deformation characterization

The volumes acquired at IOPs of 10 mmHg and 40 mmHg pre-registered well after using the pre-registration technique (Fig. 10). The calculated rigid body motion was the translation of [*T_x_, T_y_, T_z_*] = [1, 2, 5] voxels and the rotation angle of [*θ_x_, θ_y_, θ_z_*] = [-0.28°, 2.03°, 2.24°]. Fig. 11 exhibited the stretch, compression and shear strains obtained from the proposed DVC method and the conventional method. The third column shows their differences. The compression tended to be larger than the stretch. On average, the conventional DVC method estimated larger deformations than our proposed DVC method. Please recall that, by convention, compressions are negative deformations. We show them in absolute values to make things easier to compare.

**Fig. 10.**
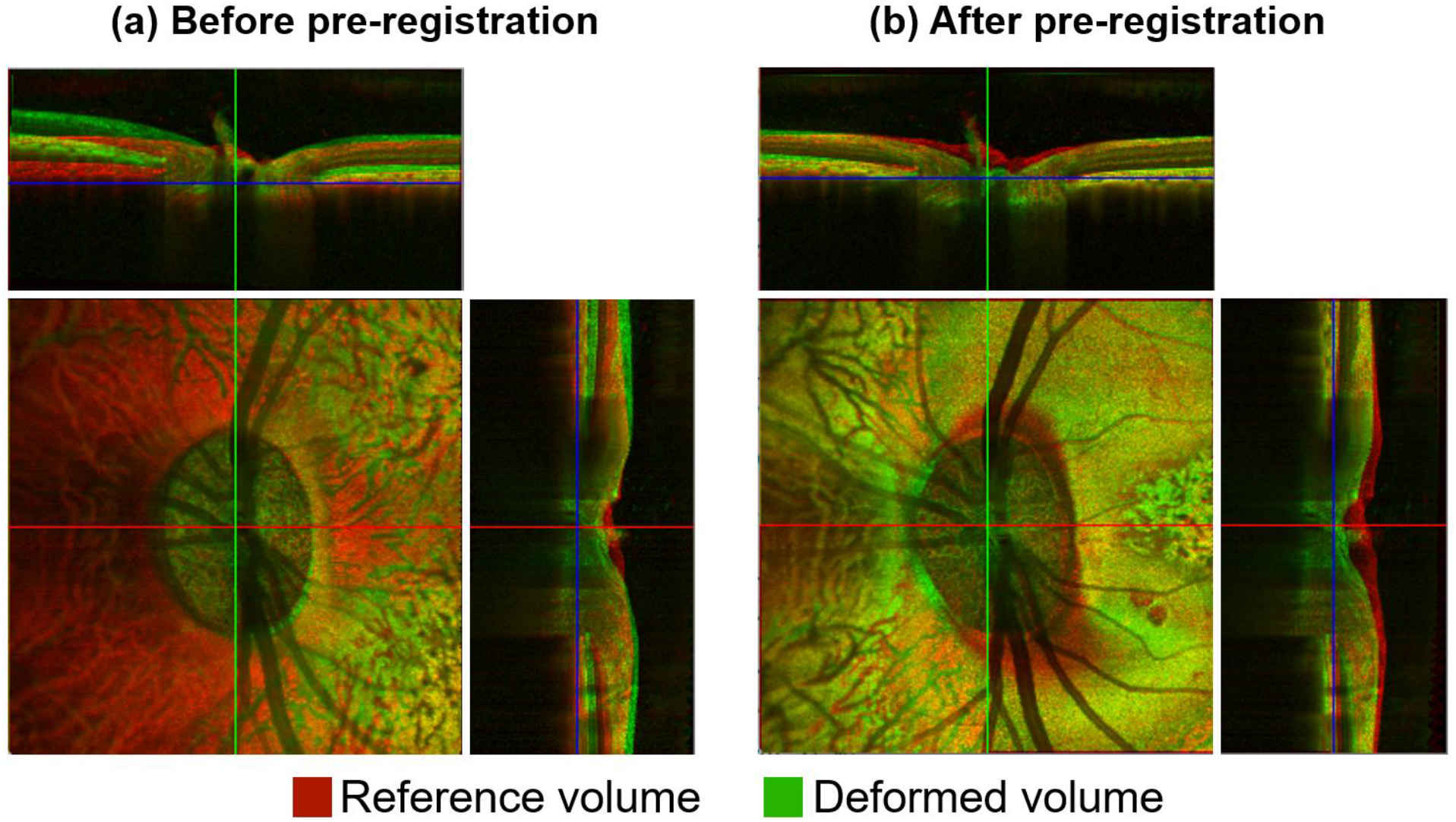
Pre-registration to correct the rigid body motion of the ONH in the OCT volumes acquired before and after elevating the IOP from 10 mmHg to 40 mmHg. (a) The mapping before the pre-registration: the red is the reference volume acquired at 10 mmHg and the green is the deformed volume acquired at 40 mmHg; (b) The mapping after the pre-registration. Note that these are actual OCT volumes, whereas Fig 7 shows artificial deformations.

**Fig. 11.**
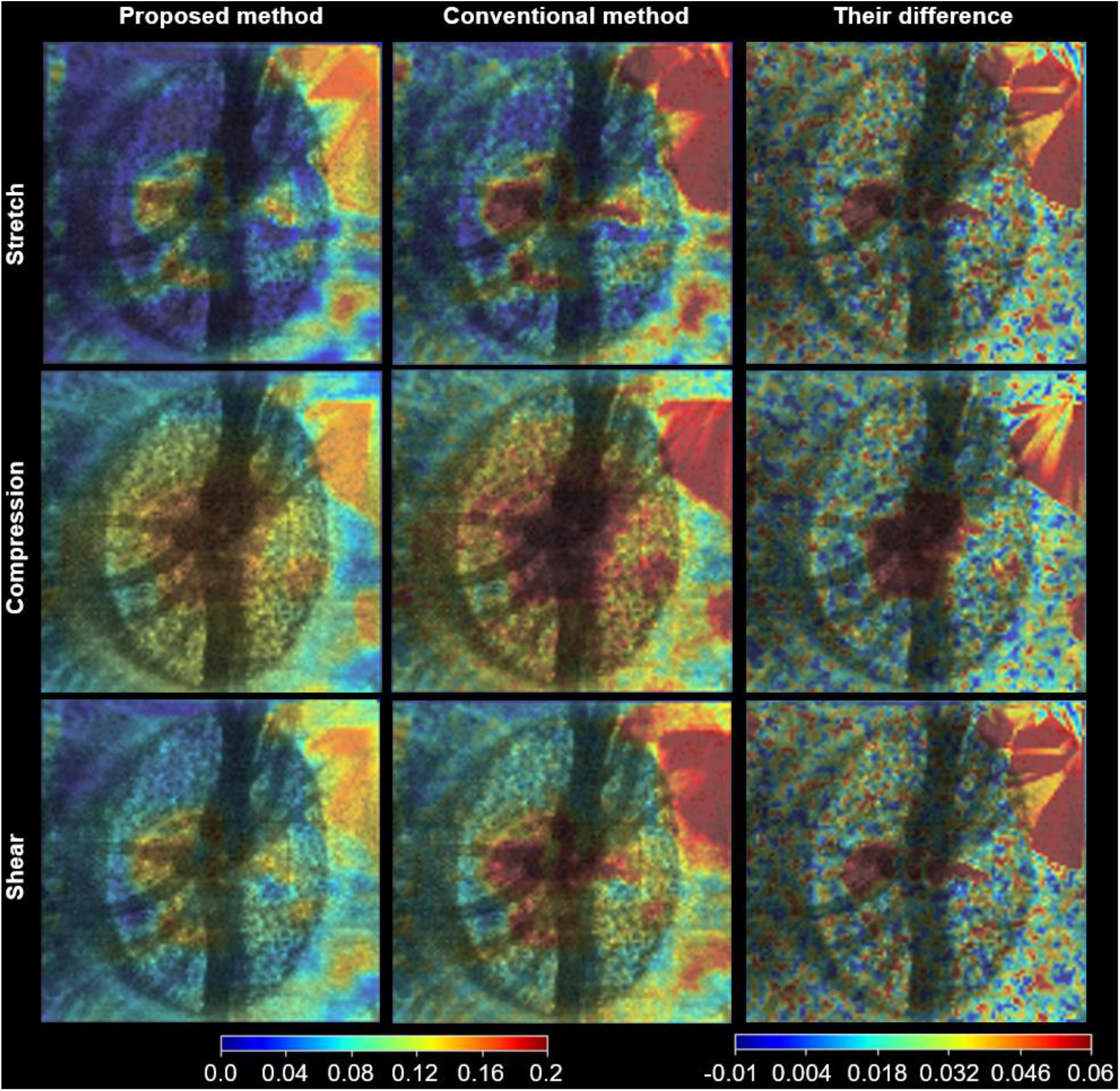
The full-field stretch, compression, and shear strain of an example C-mode of the OCT volume at the level of the lamina cribrosa induced by elevated IOP from 10 mmHg to 40 mmHg. The results in the first column were obtained with the proposed DVC method; the results in the second column were derived with the conventional DVC method; the last column shows their differences: the second column minus the first column (note smaller range spanning both positive and negative). The full-field strains were obtained from the interpolation of the calculated discrete ones. The stretch and compression are the maximal and minimal principal strains, respectively. Note that compressions are negative strains, but to simplify comparison the results are shown using the absolute values. It is observed that the compression tends to be larger than stretch and shear; on average, the conventional DVC method obtained larger strains than the proposed DVC method. Their strain differences can be up to 0.06 in some regions. The strains from the proposed DVC method should be more accurate than those from the conventional DVC method considering the fact that 1) strain calculation is very sensitive to noise and the noise often exaggerates the calculated strains; and 2) the proposed DVC method achieved more accurate strains than the conventional method on the test of artificial volumes with preset displacements and strains. From the results of the proposed DVC method, we detected larger strains (up to 0.14) in central lamina and smaller (under 0.04). Notice how the proposed method handled the complex low-signal, high-noise central LC region better than the conventional method for which the local strains become very large, suggesting an error. Both methods can show very large strains in regions where the signal quality was extremely low, for example, outside the canal in the top-right sector. These are an example of a region where it is important to consider both the strength of the correlations and the signal quality before deriving conclusions.

## 4. Discussion

Accurate characterization of ONH deformations between OCT volumes depends upon high-quality DVC. However, due to challenges in image registration, high noise, unclear accuracy, and considerable computational burden, existing DVC methods often fall short. Our goal was to introduce a revised DVC method that addresses the shortcomings of existing DVC methods through a combination of technical improvements. Four main contributions arise from this work: **1)** image pre-registration corrected ONH rigid body motion well; **2)** sub-voxel registration was achieved via a modified 3D IC-GN method, **3)** the computational burden was dramatically reduced through use of a custom look-up table and occupied memory was saved, and **4)** our DVC method had excellent overall performance in displacement and strain calculation and computation time. The synthesis of these contributions resulted in improved DVC measurement quality and workflow efficiency in real-world application. Below we discuss each of these in more detail.

### Contribution 1: Image pre-registration corrected ONH rigid body motion well

No two OCT scans are of precisely the same region in the same orientation, even when the subject is anesthetized. Eye-tracking software of some OCT systems can help reduce motion, but often there are still differences in the location or orientation of the scanned region, for example by breath or pulse, sometimes resulting in considerable ONH rigid body motion between the reference and deformed volumes. If substantial rigid body rotations remain, it could lead to failure of the DVC algorithm. Large displacements reduce computational efficiency. Reported DVC methods do not mention efforts to account for the issue of rigid body motion [7, 30, 33, 42].

We have shown that our semi-automatic pre-registration technique works well to correct ONH rigid body motion. When the ONH has a large rotation (more than 4°) between the reference and deformed volumes, manual operation was necessary; otherwise, the automatic correction would fail. Manual operation is easy and convenient due to the relatively large rotation angle error tolerance (about 2°), the developed user-friendly interactive software and the distinct edge features in the ONH, like the Bruch’s membrane. Phase correlation and Nelder-Mead nonlinear optimization were then used to further automatic calculate the rigid body motion. The deformed volume registered well to the reference volume after pre-registration. The absolute rotation angle differences (Δ*θ_x_*, Δ*θ_y_*, Δ*θ_z_*) were all less than 0.4°. This technique has the voxel-level translation accuracy, which can explain why all the absolute translation differences (Δ*T_x_*, Δ*T_y_*, Δ*T_z_*) were 0.3 voxels in the tests. ONH rigid body motions in seven actual examples were all corrected well, indicating the effectiveness of the pre-registration technique.

### Contribution 2: Sub-voxel registration was achieved via a modified 3D IC-GN method

After image pre-registration minimized rigid body motion, a coarse search was done first to locate the corresponding point with voxel level accuracy. Bar-Kochba et al. [42] implemented a coarse search process of DVC in the Fourier frequency domain as it can greatly enhance the search efficiency. However, search accuracy decreases considerably if the deformation is larger than 7%. In practice, the ONH may be subjected to deformations of more than 10% in response to elevated IOP [7]. Hence, our coarse search process was performed in the spatial domain to ensure accuracy despite its low efficiency. Then, the popular 3D IC-GN iteration method was used to obtain the deformation parameter p including displacement vector of sub-voxel accuracy by minimizing the ZNSSD coefficient. Girard et al. [7] directly applied a genetic optimization algorithm to determine the deformation gradient tensor and displacement vector by minimizing the ZNSSD coefficient. This method can achieve high accuracy, but its computational efficiency is very low, i.e., 15 hours to process 10,000 POIs. Compared with the genetic optimization algorithm which was directly applied to determine the deformation gradient tensor and displacement vector by minimizing the ZNSSD coefficient [7], 3D IC-GN has substantially higher computation efficiency, i.e., less than 18 minutes to process 27, 000 POIs. OCT images can have low contrast and considerable noise. Even when the OCT instrument’s signal averaging function is used, the acquired volumes in our practical applications still have relatively high noise level. This issue hinders the convergence of 3D IC-GN iteration. When it fails to converge, we selected the p at the iteration step having the minimal ∥Δ*p*∥ instead of the minimal *C_ZNSSD_* in order to ensure sub-voxel accuracy of DVC. ∥Δ*p*∥ was likely to be at its minimum when p most approaches the exact value in the iteration process, whereas, the difference of *C*_□□□□□_ is negligible, which can also explain why the convergence condition is set based on ∥Δ*p*∥ instead of *C*_□□□□□_. With the help of the modified 3D IC-GN method for sub-voxel registration, our proposed DVC method had excellent accuracy in sub-voxel translation and strained deformation measurement. The average absolute displacement errors in the X, Y, and Z directions were very similar, under 0.028 voxels with speckle noise and under 0.037 voxels with Gaussian noise, indicating its isotropic accuracy in the three directions. The respective average, absolute, and relative strain errors in these three directions were less than 0.0018 and 4% with speckle noise, and less than 0.0045 and 8% with Gaussian noise. Gaussian noise had a more negative effect on DVC accuracy than the same level of speckle noise. In fact, speckle noise of the actual OCT volume was substantially larger than Gaussian noise. As a result, the former practically results in greater DVC measurement error than the latter.

### Contribution 3: Computational burden of non-integer voxel interpolation was reduced by a custom look-up table and occupied memory was saved

Conventional DVC is computation- and time-intensive partly because of a large number of interpolations which are required with sub-voxel registration to calculate non-integer voxel intensities. In each iteration, the number of interpolations is equal to the size of subvolume, up to 12,000 non-integer voxel interpolations in this work. In other studies, a trilinear interpolation method was employed [7, 42] due to its high computational efficiency and ease of implementation. Yet, its interpolation error is not negligible. The popular higher-order tricubic B-spline interpolation method was used in this DVC method because of its reduced interpolation error, but its main shortcoming has very low computational efficiency. To speed up computation, we built a look-up table to save the results of the first three multiplicative factors of Eq. (8). This look-up table was independent of the processed OCT volume and only occupied 0.61 MB. Each non-integer voxel interpolation in our method consisted of 84 multiplicative and 63 additive calculations. Optimized computation dramatically increased DVC efficiency by more than 50% and saved the memory by about 30%, comparing with the conventional method using a look-up table of B-spline control points (more than 800 MB memory, 212 multiplicative and 155 additive calculations) [33] and conventional method using a look-up table of 64 interpolation coefficient (more than 50 GB, 192 multiplicative and 63 additive calculations) [30]. If higher resolution OCT volumes are used, the proposed interpolation method would have a more direct advantage as its look-up table is independent of the volume and remains the same, whereas, the conventional methods would consume more memory.

### Contribution 4: Our DVC method had excellent overall performance in displacement and strain calculation and computation time

With rigid body motion, the overall performance of our DVC method had fairly considerable advantages over the conventional DVC method in displacement and strain measurement accuracy, and computation efficiency. Our DVC method had overall displacement errors smaller than 0.05 voxel with speckle noise and 0.07 voxel with gaussian noise, and the overall strain errors of under 0.0025 with speckle noise and 0.006 with Gaussian noise. Overall displacement and strain errors of our DVC method were less than 1/3 of the conventional method. Besides, our DVC method takes less than 18 minutes to process 27 000 POIs; the computation efficiency was enhanced by about 75%. If the rotation angle is relatively large, i.e., up to 7 degrees, the conventional DVC method would fail to work, whereas, it would not affect the proposed DVC method because of the presented pre-registration technique. The excellent performance of our DVC method was also partly the result of the confidence-guided searching strategy and confidence weighted strain calculation: 1) the number of low reliable POIs misused to guide the searching path in our method was decreased to under 1/3 of that of conventional correlation-coefficient-guided searching strategy; and the strain obtained from the confidence weighted strain calculation method was also more accurate than the conventional strain calculation method. The confidence was defined by considering both image contrast of subvolume and correlation coefficient.

### Synthesis in application

In application, our DVC method resulted in high quality measurements of ONH deformation. After using the described pre-registration technique, ONH images acquired at the elevated IOP of 40 mmHg registered very well to their reference volumes acquired at 10 mmHg. The measured ONH deformations were complex, especially in the regions of central laminar cribrosa and sometimes on the periphery. The maximal stretch, compression and shear strains did not always colocalize. There were relatively large differences (up to 0.06) of the measured stretches, compressions, and shears between the proposed DVC method and the conventional DVC method. The conventional DVC method tended to overestimate the ONH deformation because strain calculation is very sensitive to noise which often exaggerates the calculated strains.

### Recommendations

Despite relatively large amplitude noise, we recommend against the use of OCT image pre-processing with this DVC method except for an image median filter. In particular, image contrast enhancement and histogram normalization should be avoided. These operations are useful for better visualization of OCT volumes, but they change the real voxel intensity variation information which may negatively affect the DVC accuracy. In our tests, we did not apply any image pre-processing operations to the OCT volumes in order to better reflect the performance of the DVC method. We did not test for the potential effects of signal compensation techniques [43, 44], although our results suggest that these are not necessary. Additionally, a raster, instead of radial, scan model is recommended for ONH imaging using the OCT because the former allows sampling with the same resolution over the whole volume, leading to consistent DVC measurement accuracy of the whole volume, whereas, the latter reduces in resolution with the distance from the scan’s center.

### Remaining challenges and considerations

There are some aspects of the proposed DVC method that deserve further consideration. The shadow of the blood vessels in the OCT volume is a challenging issue affecting the measurement accuracy of the POI near the shadow. Several previous studies have used compensation or similar techniques to reduce the shadows [43, 44]. It remains unclear how much the compensation may affect DVC-measured displacements and deformations within the shadows and deep within the tissues. We opted instead for identifying the shadow regions and removing them from analysis to attenuate their influence. Image segmentation along with morphological analysis, such as erosion and dilation, was adopted to separate the shadow from the ONH in this work. Nevertheless, it is difficult to choose the optimal threshold for segmentation: if the threshold is too large, some important regions may be missed; if it is too small, the shadow region is difficult to remove completely. It may be possible to improve this process by using deep learning methods to separate out the shadows or use the OCT-angiography information from the same scans to predict the location of the most problematic shadows. Setting a relatively large size of the subvolume for correlation is also useful to minimize the influence of shadows. Another consideration is the manual component of our semi-automatic pre-registration technique. Effective manual operation more or less depends on the user’s experience to identify and coarsely align the ONH’s fairly apparent key features. To achieve fully automatic pre-registration, we may apply 3D scale-invariant feature transform (SIFT) [45] feature points to align the ONH in the future. Lastly, we can perform the DVC technique in the graphics processing unit (GPU) platform to further improve its computational efficiency, although the proposed method has sped up the computation greatly compared with the conventional DVC method.

### Summary

We present a high-accuracy and high-efficiency DVC technique to characterize in-vivo ONH deformations from OCT volumes. The method has been successfully applied to characterize the deformation of monkey ONHs subjected to acute and chronic IOP elevation [46, 47]. This technique has the potential to help investigate the pathologic mechanism of glaucoma and eventually, to help clinically diagnose glaucoma in its early stages. Although we demonstrate efficacy of this tool for images of the ONH, this DVC method can also be used to characterize the biomechanics of other biological tissues [48–50].

