## Supplementary Material 1 for "A high-accuracy and high-efficiency digital volume correlation method to characterize *in-vivo* optic nerve head biomechanics from optical coherence tomography"

### 1 Phase correlation to further correct the rigid body motion

In order to correct the translation, we assume that there is only a translation between the reference subvolume $f(x, y, z)$ and the deformed subvolume $g_1(x, y, z)$ even if relatively small rotation angle (less than 2 degree) still exists after manual operation,

$$f(x,y,z) = g_1(x - u_0, y - v_0, z - w_0) \quad \text{(S1)}$$

Appling the fast Fourier transform (FFT) to both $f$ and $g_1$,

$$F(\alpha,\beta,\gamma) = e^{-j2\pi(\alpha u_0+\beta v_0+\gamma w_0)} G_1(\alpha,\beta,\gamma) \quad \text{(S2)}$$

The cross-power spectrum of two images $f$ and $g_1$ with Fourier transform $F$ and $G_1$ is defined as,

$$\frac{{G_1}^*(\alpha,\beta,\gamma)F(\alpha,\beta,\gamma)}{|{G_1}^*(\alpha,\beta,\gamma)||F(\alpha,\beta,\gamma)|} = e^{-j2\pi(\alpha u_0+\beta v_0+\gamma w_0)} \quad \text{(S3)}$$

where ${G_1}^*$ is the complex conjugate of $G_1$. The peak position ($u_0$, $v_0$, $w_0$) of the inverse Fourier transform of Eq. (S3) is regarded as the translation from $g_1$ to $f$.

Subsequently, phase correlation is used to rectify the rotation angles ($\theta_x$, $\theta_y$, $\theta_z$) of the deformed volume $g_2$ separately. $g_2$ is obtained from shifting $g_1$ by ($u_0$, $v_0$, $w_0$). We take the correction of the rotation angle $\theta_z$ in the Z direction as the example. Assume that there is only a rotation angle $\theta_z$ between $f$ and $g_2$,

$$f(x,y,z) = g_2(x\cos\theta_z + y\sin\theta_z, -x\sin\theta_z + y\cos\theta_z, z) \quad \text{(S4)}$$

Appling the fast Fourier transform (FFT) to both $f$ and $g_2$ and we only considered the magnitude of the spectrum,

$$|F(\alpha,\beta,\gamma)| = |G_2(\alpha\cos\theta_z + \beta\sin\theta_z, -\alpha\sin\theta_z + \beta\cos\theta_z, \gamma)| \quad \text{(S5)}$$

The magnitudes are then defined in the polar coordinates to obtain the rotation angle component:

$$|F(\rho,\theta,z)| = |G_2(\rho,\theta - \theta_z, z)| \quad \text{(S6)}$$

It can be seen that the rotation in the Cartesian coordinate can be transformed into a translation in the polar coordinate. FFT is then applied to the spectrum magnitude images $|F(\rho,\theta,z)|$ and $|G_2(\rho,\theta-\theta_z,z)|$ in polar coordinate,

$$F'(\alpha',\beta',\gamma') = {G_2}'(\alpha',\beta',\gamma')e^{-j2\pi\beta'\theta_z} \tag{S7}$$

where $F'(\alpha',\beta',\gamma')$ and ${G_2}'(\alpha',\beta',\gamma')$ are the spectrum of $|F(\rho,\theta,z)|$ and $|G_2(\rho,\theta-\theta_Z,z)|$, respectively. The rotation angle $\theta_z$ is then calculated by the inverse Fourier transform of the follow equation (the same as Eq. (S3)),

$$\frac{{G_2}'^*(\alpha',\beta',\gamma')F'(\alpha',\beta',\gamma')}{\left|{G_2}'^*(\alpha',\beta',\gamma')\right|\left|F'(\alpha',\beta',\gamma')\right|} = e^{-j2\pi\beta'\theta_z} \tag{S8}$$

where ${G_2}'^*$ is the complex conjugate of ${G_2}'$. The rotation angles $\theta_x$ and $\theta_y$ are then calculated in the same way as Eqs. (S4)-(S8). Subsequently, a nonlinear optimization method, Nelder-Mead method, is further used to optimize ($\theta_x$, $\theta_y$, $\theta_z$) by minimizing the $[f(x,y,z) - g_2(x^*,y^*,z^*)]^2$, where

$$\begin{bmatrix} x \\ y \\ z \end{bmatrix} = \begin{bmatrix} \cos\theta_z & \sin\theta_z & 0 \\ -\sin\theta_z & \cos\theta_z & 0 \\ 0 & 0 & 1 \end{bmatrix} \begin{bmatrix} \cos\theta_y & 0 & \sin\theta_y \\ 0 & 1 & 0 \\ -\sin\theta_y & 0 & \cos\theta_y \end{bmatrix} \begin{bmatrix} 1 & 0 & 0 \\ 0 & \cos\theta_x & \sin\theta_x \\ 0 & -\sin\theta_x & \cos\theta_x \end{bmatrix} \begin{bmatrix} x^* \\ y^* \\ z^* \end{bmatrix} \tag{S9}$$

In practice, we can also start the next round phase correlation to rectify the rigid-body motion again until satisfactory results. It is noted that 1 or 2 round phase correlation is enough for most situations.

### 2 Multi-thread parallel computation

The computation intensity of the DVC is very high because the size of the subvolume for correlation is often more than 10 000 voxels and a large number of POIs is required to be searched. Hence, we try to speed up the computation using the multi-thread parallel computation technique. At first, we select 1 POI at the center of region of interest (ROI) and another 6 candidate POIs around that central one. The searching area for the first POI is relatively large since we do not know how large the deformation is. Multi-thread computation technique is used to simultaneously calculate the correlation coefficients between the reference subvolume centred at the first POI and multiple deformed subvolumes. It is noted that if the confidence *Conf* of this first POI is lesser than the threshold $T_{conf}$ (set as 0.70 in this work), we will choose another candidate POI as the first POI till its *Conf* is larger than $T_{conf}$. Subsequently, the surrounding unsearched POIs of the POI with the *Conf* higher than $T_{conf}$ are added into the queue of the searching list. Multi-thread computation technique is then used to simultaneously find the corresponding points for multiple POIs in the queue of the waiting list. The positions of the corresponding points of the unsearched POIs in the queue are estimated by the positions of the reliable neighboring searched POI. The majority of POIs are searched when the searching list is cleared. However, there are still some POIs missed to be searched if the *Conf* of its neighboring POIs are all lesser than $T_{conf}$. The remaining unsearched POIs are also simultaneously searched using multi-thread parallel computation according to the average displacement of the searched POIs. Fig. S1 shows the workflow of the proposed DVC using the multi-thread parallel computation technique.

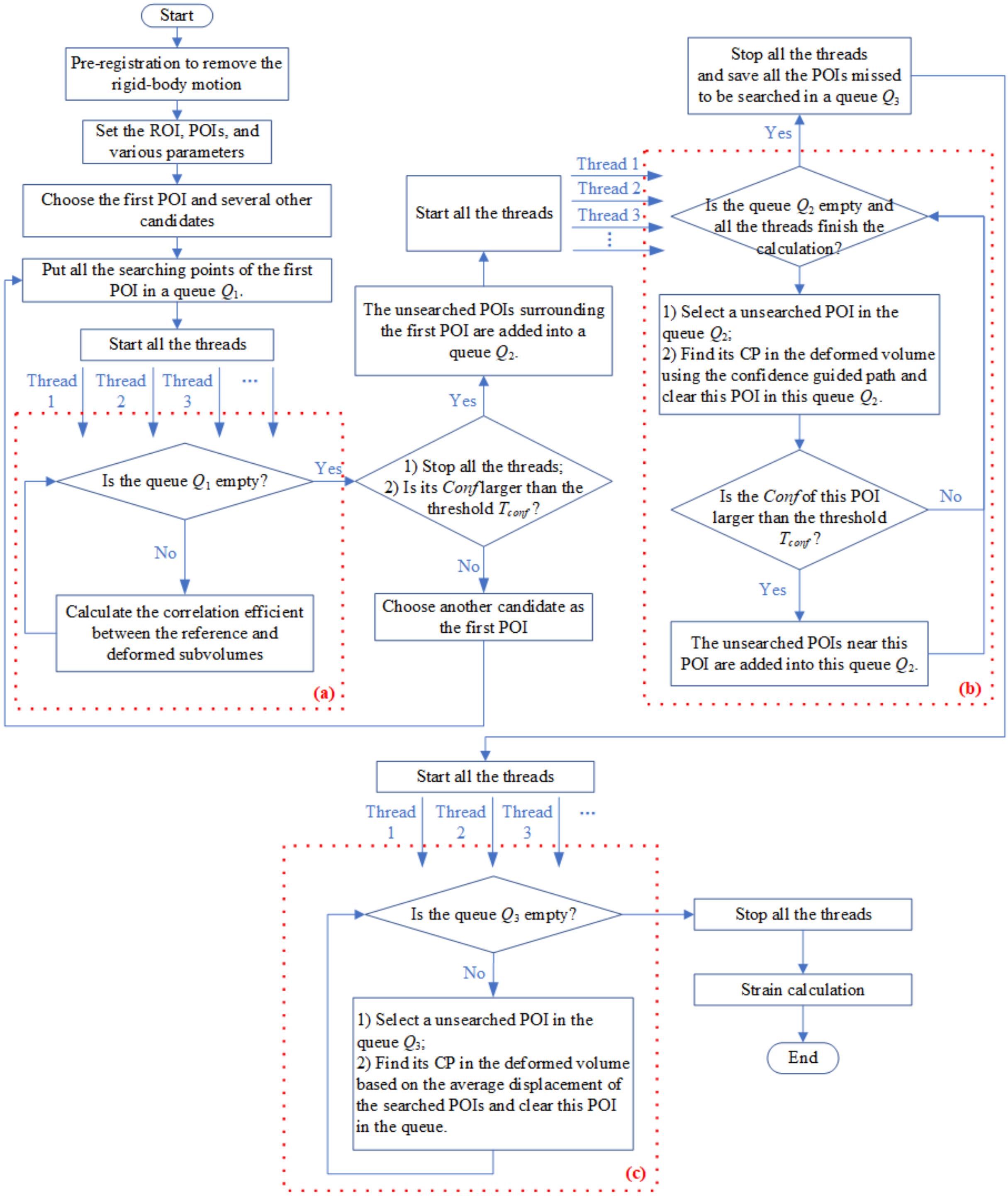


**Fig. S1**. The workflow of the proposed DVC method. The multi-thread parallel computation is applied to the parts (a), (b) and (c).
