## Supplementary Material 2 for "A high-accuracy and high-efficiency digital volume correlation method to characterize *in-vivo* optic nerve head biomechanics from optical coherence tomography"

The presented pre-registration technique is tested on 7 pairs of real OCT volumes acquired during the chronic remodeling of the ONH. The results show the (green) deformed volumes register the (red) reference volumes well.

**Before pre-registration** **After Pre-registration**

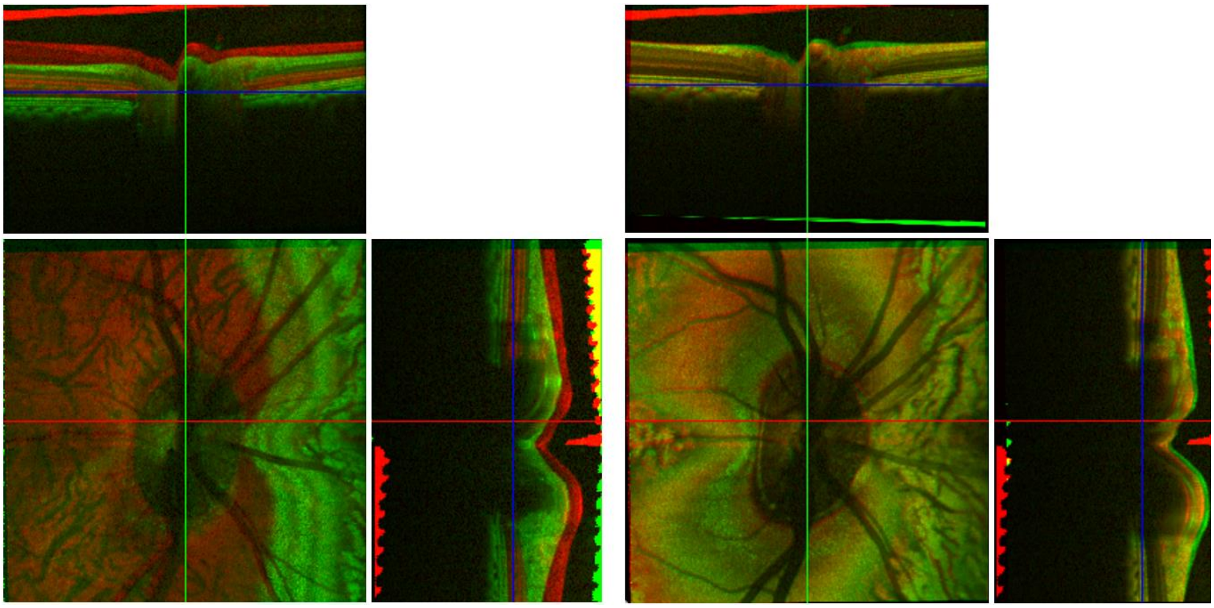

**(1)**

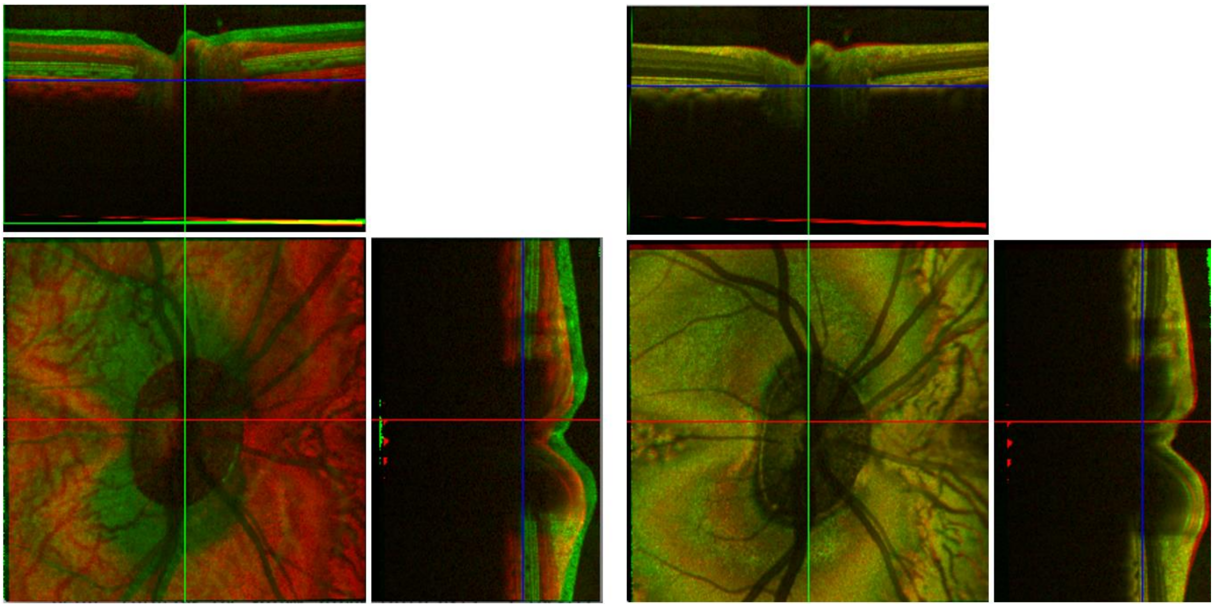

(2)

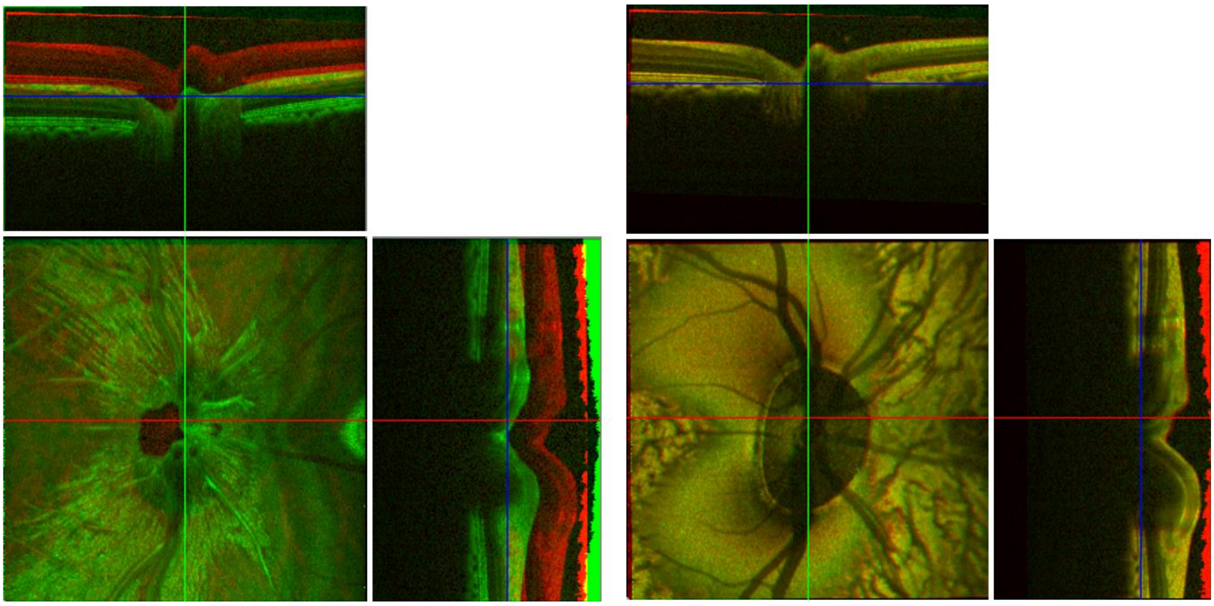

**(3)**

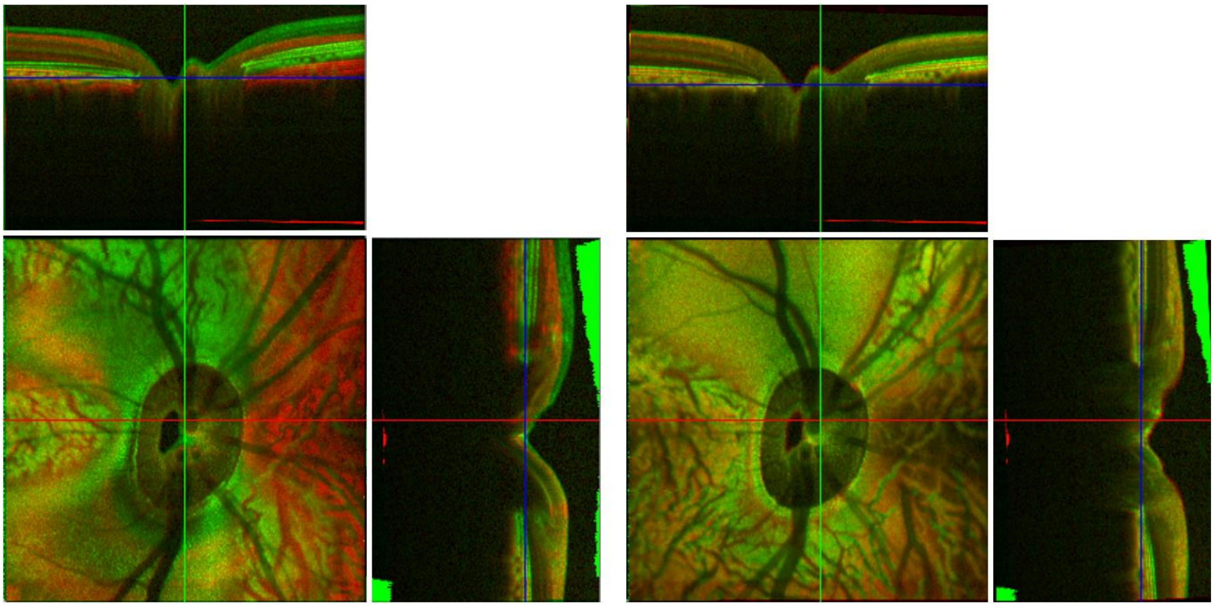

**(4)**

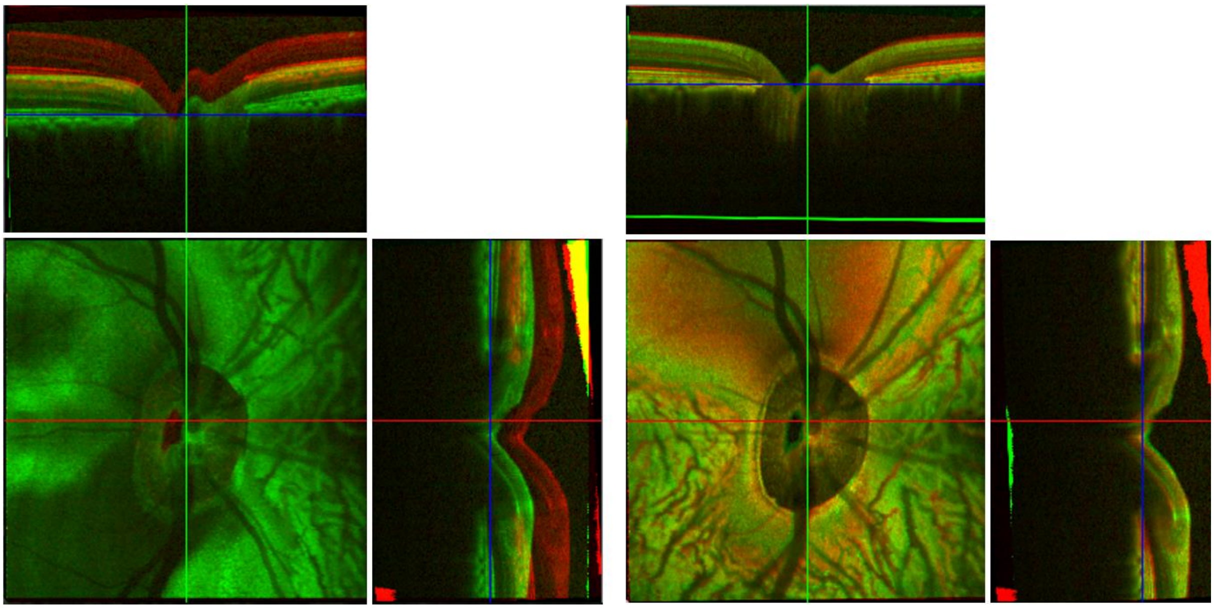

**(5)**

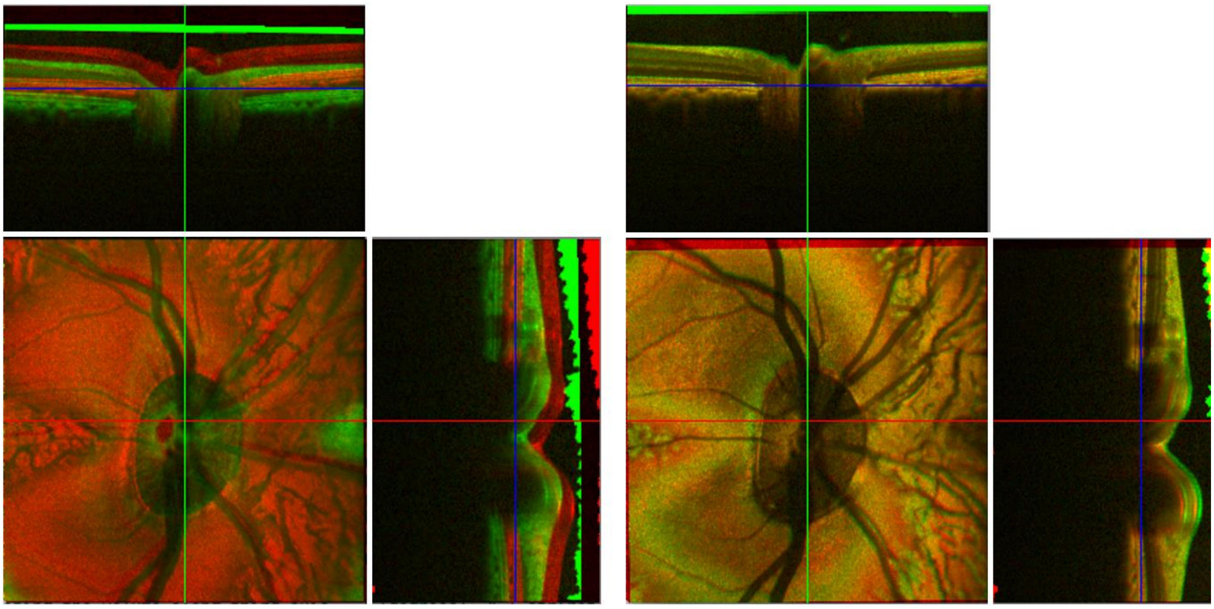

**(6)**

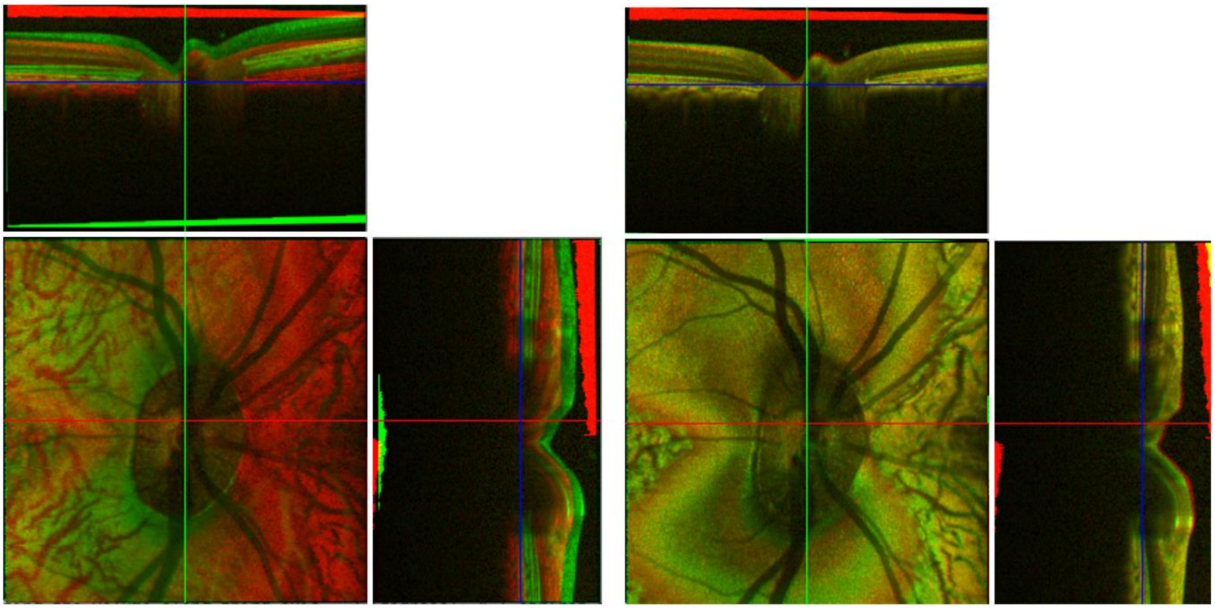

(7)
