## Supplementary Material 3 for "A high-accuracy and high-efficiency digital volume correlation method to characterize *in-vivo* optic nerve head biomechanics from optical coherence tomography"

### 1. Computation efficiency evaluation

The proposed DVC method consists of three parts: 1) pre-registration to remove the rigid-body motion of the ONH; 2) DVC method to track the corresponding points between the reference and deformed volumes, including coarse searching and 3D IC-GN sub-voxel registration; 3) strain calculation from the calculated displacements. Among the three steps, step 2 has the highest computation intensity and occupies the majority computation time. In step 1, once manual operation is done to register the deformed volume to the reference volume according to the edge information, automatic pre-registration takes less than 1 minute to further correct the rigid-body motion. The strain calculation time in step 3 is almost negligible, relative to the computation time in step 1 and 2. The parameters of the used computer are: Intel Core i7-8750H CPU @2.20GHz and RAM 16.0 GB. The operation system is Windows 10 and the programming platform is Visual Studio 2019. Our code is performed using 11 computation threads. Once the hardware is determined, the computation time of corresponding point tracking in step 2 depends on the number of POIs, size of subvolume, searching area, and the deformation degree. The number of POIs is 26818 and the size of subvolume is 23×23×23. In the practical application, the deformation is often not known at first, so we have to choose a relatively large searching area. In this test, we select the same searching area for all the OCT volumes: the searching areas for the first batch POIs, path-dependent POIs, and the missed searched POIs are 25×25×25, 3×3×3, and 13×13×13, respectively. Fig. S1 shows the average computation time of 5-round repeated corresponding point tracking in step 2. It can be seen that the computation time under rigid translation is slightly more than 8 minutes; while the computation time under strained deformation increases from about 9 minutes to about 17 minutes with the increment of the strained deformation. The computation time in the situation of Gaussian noise is slightly higher than that in the situation of speckle noise since the 3D IC-GN iteration number in the former is more than that in the latter. The computation time under rigid translation only is relatively stable and less than that under strained deformation. The reason why the computation time increases with the increment of the strained deformation is that the increased deformation would reduce the similarity between the reference subvolume and the target subvolume, thereby increasing the number of missed searched POIs during the path-dependent searching process. The searching area of the missed searched POIs is much larger than that of the path-dependent searched POIs. In addition, the large deformation also increases the 3D IC-GN iteration number before the convergence.

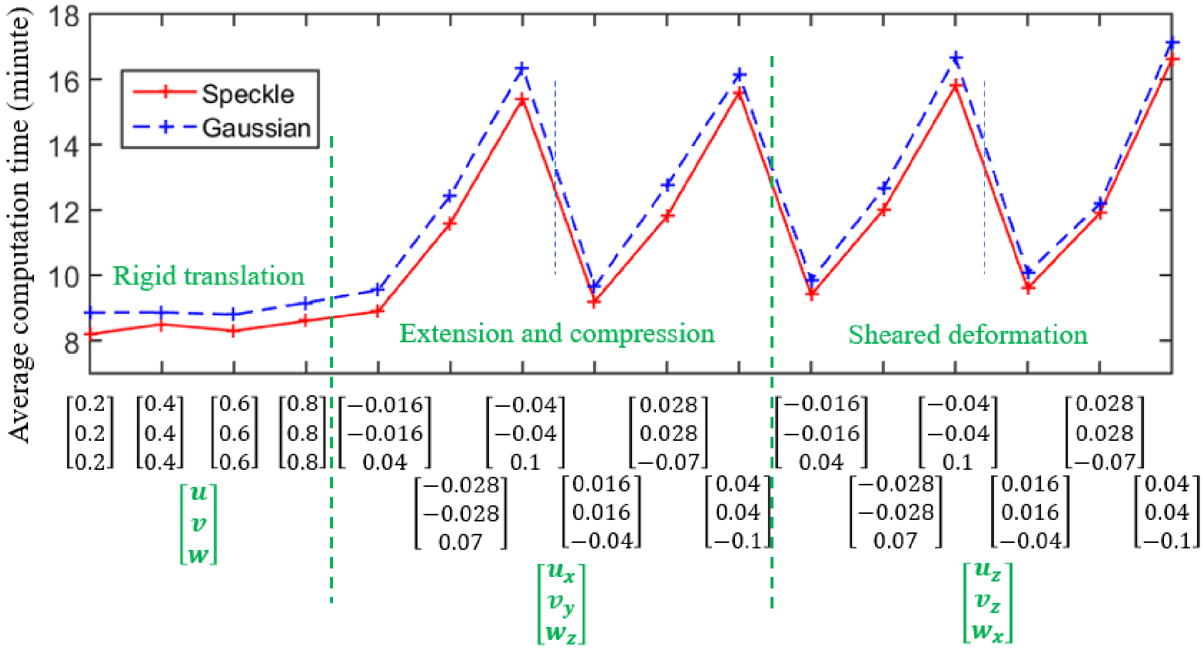


**Fig. S1.** The average computation time of 5-round repeated corresponding point tracking under rigid translation, extension, compression and sheared deformation.

### 2. The proposed sub-voxel registration methods (Method 1 and Method 2) *vs* the conventional sub-voxel registration method

Sub-voxel registration is necessary to guarantee the accuracy of the corresponding point. However, the weak texture, low contrast, but high noise level of the OCT volume of the ONH lows down the convergence ability of the 3D IC-GN iteration. In the practical application, more than 90% of the POIs exceed the iteration number limit and fail to converge if the convergence criterion is set as $||\Delta\mathbf{p}|| \leq$ 0.001 which is the most common criterion; if we low down the convergence criterion to 0.01, about 34% of POIs still fails to converge; if we further low down the convergence criterion to 0.1, although the majority POIs can converge, the accuracy of the tracked corresponding point may not be enough. Hence, the convergence criterion is recommended as 0.01 or 0.001, but a lot of POIs fail to converge. In the conventional method, the initial point having the lowest $C_{ZNSSD}$ from the coarse searching process is commonly selected as the corresponding point of the POI if it fails to converge, but it only has voxel-level accuracy. In this work, we still need to find the sub-voxel-accuracy corresponding point even if the POI fails to converge. When the iteration number exceeds the limit, we deduce the sub-voxel-accuracy corresponding point using the **p** having 1) the minimal $||\Delta\mathbf{p}||$ (Method 1) or 2) the minimal $C_{ZNSSD}$ (Method 2). We compare the overall displacement errors ($\Delta d = \sqrt{\Delta u^2 + \Delta v^2 + \Delta w^2}$) of the Method 1, Method 2, and the conventional method in the situation of speckle noise and Gaussian noise under the strained deformation: $[u_x, v_y, w_z]$ = [-0.04, -0.04, 0.1] and [0.04, 0.04, -0.1]; $[u_z, v_z, w_x]$ = [-0.04, -0.04, 0.1] and [0.04, 0.04, -0.1]. Fig. S2 shows the result. If the convergence criterion is set as 0.1, the displacement errors of the three methods are almost the same because all the POIs can converge. When the convergence criterion is set as 0.01 or 0.001, Method 1 has slightly higher accuracy than Method 2, while the accuracy of the conventional method is the lowest. When the convergence criterion is 0.001, the accuracy of the conventional method is low since about 71% of the POIs which fail to converge only has voxel-level-accuracy corresponding points. The displacement accuracy in the situation of speckle noise is smaller than that in the situation of Gaussian noise, which agrees with the result in the subsection. In addition, the displacement accuracy is only enhanced slightly if the convergence criterion reduces from 0.01 and 0.001, however, the iteration time is increased by about 78%. Hence, the convergence criterion is recommended as 0.01 to balance the accuracy and efficiency.

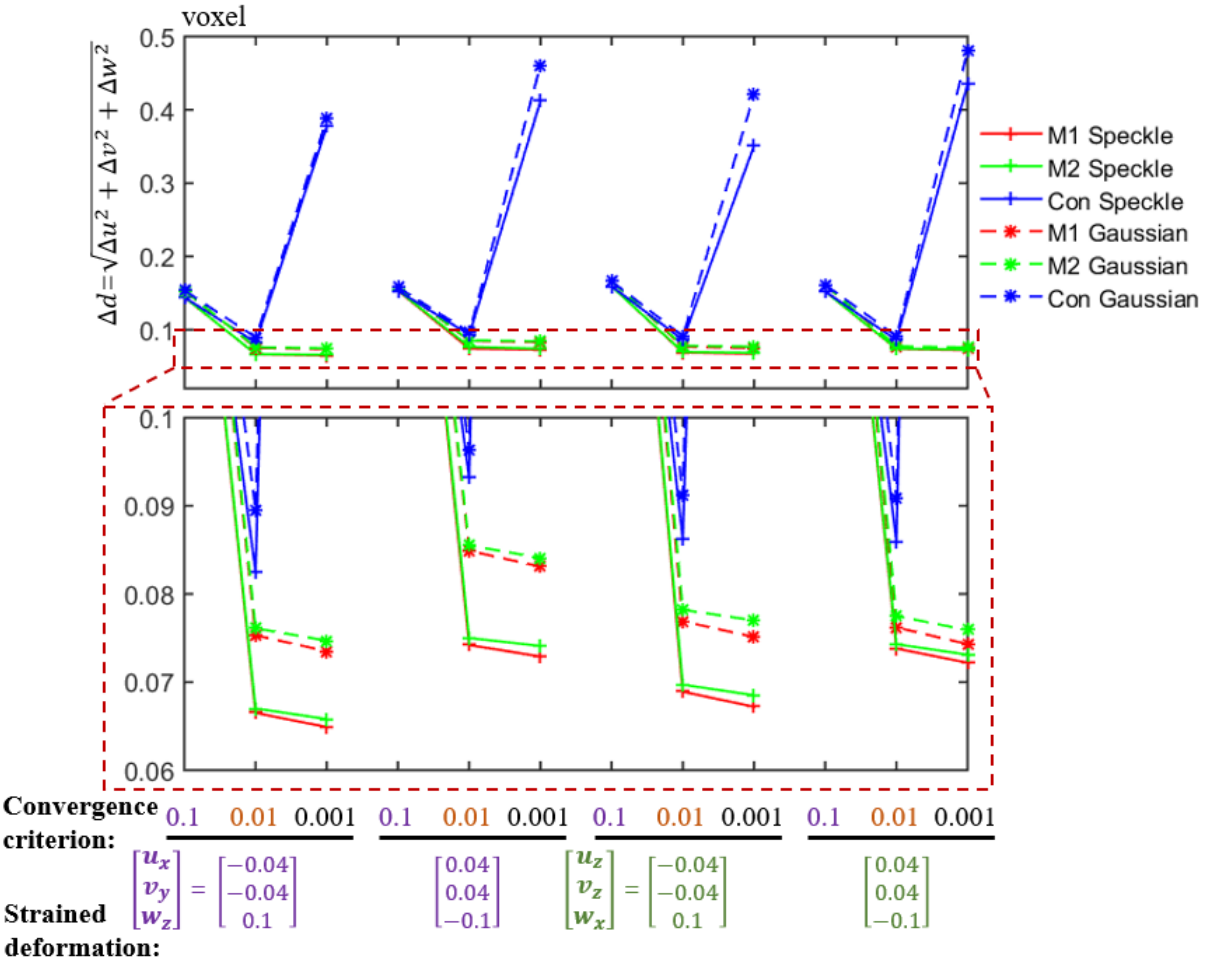


**Fig. S2.** The overall displacement error $\Delta d = \sqrt{\Delta u^2 + \Delta v^2 + \Delta w^2}$ of the sub-voxel registration using the **p** having 1) the minimal $||\Delta \mathbf{p}||$ (Method 1: **M1**) and the minimal $C_{ZNSSD}$ (Method 2: **M2**), and the conventional method (**Con**) under various strained deformations in the case of speckle noise and Gaussian noise.

#### 3. The proposed fast and memory saving tricubic B-spline interpolation method vs the existing methods

In the 3D IC-GN iteration method, there is a large number of interpolation calculations, i.e., in each iteration, we need to interpolate $(2M+1)^3$ voxels which is the size of the subvolume for correlation. For example, in this study, 23×23×23 = 12167 voxels are required to be interpolated in each iteration. Trilinear interpolation is very efficient, but its accuracy is not enough. Tricubic B-spline interpolation method is very popular because of its higher accuracy than the trilinear interpolation method. However, its computation intensity is very high. In order to speed up the computation, one method is to build a look-up table for all the interpolation coefficients of each voxel [30] and the interpolated voxel intensity can then be directly calculated by

$$\mathrm{g}(x,y,z)=\sum_{i=0}^{3}\sum_{j=0}^{3}\sum_{k=0}^{3}\alpha_{ijk}(x-[x])^{i}\,(y-[y])^{j}(z-[z])^{k} \tag{S1}$$

where $\alpha_{ijk}$ represent the 64 interpolation coefficients saved in the look-up table. Eq. (S1) is involved with about 192 multiply calculations and 63 additive calculations. The main drawback of this method is the large consumption of the memory. For example, if each data is saved as the *float* type, in this work, they consume more than 50 GB memory, which is not feasible. Additionally, the calculation of the 64 interpolation coefficients for each voxel of the OCT volume in advance is also complex and time consuming. Another method is to build a look-up table to save the control points ***R*** for each layer of the OCT volume in advance [32], it can reduce the redundant calculation and consumes much less memory than the method does in Eq. (8), for example, it occupies about 800 MB memory to save the control points ***R*** for the OCT volume used in this work. However, each non-integer voxel interpolation in this method includes 212 multiply calculations and 156 additive calculations. This method also has the same limitation as the first method: once the deformed OCT volume is changed, we need to update the look-up table to save the control points which is also a complex process. In the proposed method, we only need to build up a look-up table for the results of the first three multiply factor in Eq. (6): *LUT* = $[1\ \ t-[t]\ \ (t-[t])^2\ \ (t-[t])^3]\,\boldsymbol{K}\,(\boldsymbol{A}_{4\times4})^{-1}$. This look-up table is independent to the processed OCT volume, indicating that there is no need to update the table when the deformed OCT volume is changed, which is an advantage over the two existing methods. In this work, $t-[t]$ ranges from 0 to 1 at the step of 0.00005, this table only consumes about 0.61 MB memory if the data is saved as *double* type, much less than the two existing methods. Moreover, each non-integer voxel interpolation contains 84 multiply calculations and 63 additive calculations, much less computation than the two existing methods. Table S1 shows the comparison between the proposed method and the two existing methods. It demonstrates the obvious advantage of proposed method over the two existing methods in terms of computation efficiency and memory consuming. The two existing methods should have the same accuracy, whereas, the method to establish a look-up table of 64 interpolation coefficients occupies too much memory. Hence, we only compare the accuracy and computation time of the proposed method with that of the method to build the look-up table of the control points under various strained deformation in the situation of Gaussian noise and speckle noise.

**Table S1**. The comparison between the proposed tricubic B-spline interpolation method and the two existing methods.

| Methods / Properties | Look-up table of 64 interpolation coefficients | Look-up table of the control points | Proposed method |
|---|---|---|---|
| Calculation intensity of each sub-voxel interpolation | 192 multiply + 63 additive calculations | 212 multiply + 156 additive calculations | 84 multiply + 63 additive calculations |
| Memory consuming | > 50 GB | > 800 MB | about 0.61 MB |
| Establishing of the look-up table | Complex, dependent to the deformed volume | Complex, dependent to the deformed volume | Simple, invariant to the deformed volume |

It is noted that the occupied memory is estimated based on the size of the processed OCT volume of an ONH: 768×769×380 voxels.

Fig. S3 shows the 3D IC-GN computation time and the overall displacement error of the proposed method and the existing method. It is noted the computation time of establishing the look-up table is also considered. The computation time of the existing method is more than 2 times to that of the proposed method, whereas, the accuracy of the former is only slightly higher than that of the latter, nearly negligible.

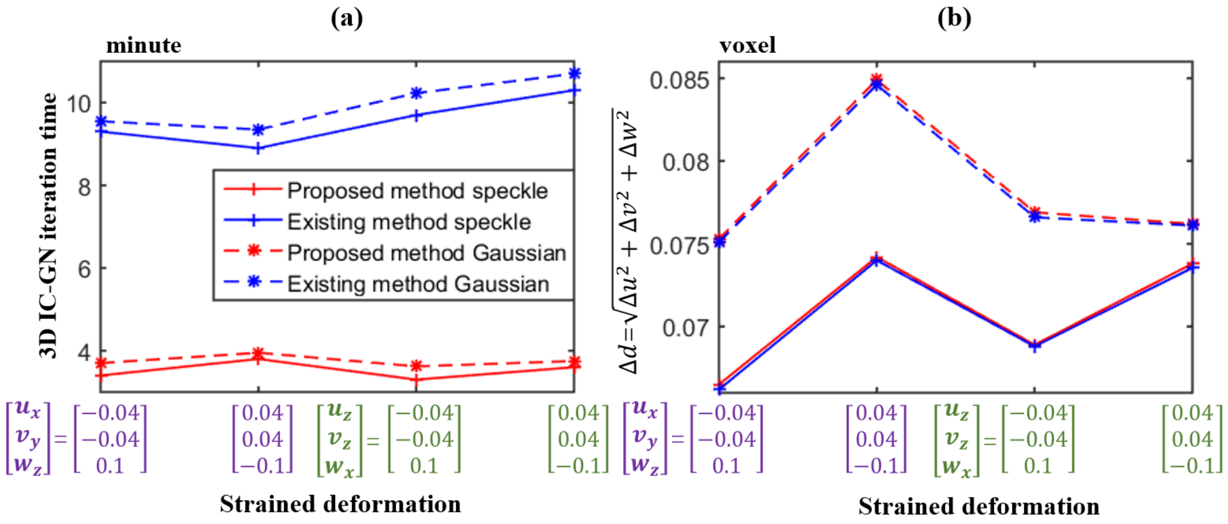


**Fig. S3.** The computation time of the 3D IC-GN iteration (a) and the overall displacement error ($\Delta d = \sqrt{\Delta u^2 + \Delta v^2 + \Delta w^2}$) (b) of the proposed tricubic B-spline interpolation method and the existing method to build the look-up table of the control points.

### 4. The proposed confidence-guided searching strategy vs the conventional correlation-coefficient-guided searching strategy

In the corresponding point searching process of the DVC, conventionally, the correlation coefficient *Corr* is often used to guide the searching path by propagating the deformation parameters from the searched POI with high *Corr* to the neighboring uncalculated POIs and estimating the positions of their corresponding points. Because of the weak texture and low contrast of the OCT volume of the ONH, sometimes, the high *Corr* of the POI does not mean its high reliability. If the low reliable POIs are misused to estimate the corresponding points of the neighboring unsearched POIs and pass the deformation parameters, the computation robustness of the DVC will be negatively affected. In this work, we combine *Corr* with AVIG to define the confidence *Conf* of the POI, expressed by the Eqs. (9) and (10), where AVIG can indicate the contrast of the subvolume centered at the POI. The *Conf* is then used to guide the searching path of the proposed method. We compare the proposed method to the conventional method in terms of the number of low reliable POIs misused to guide the searching path. The result is shown in Fig. S4. That number of conventional *Corr* guided method has more than three times to that of the proposed *Conf* guided method, indicating the much higher computation robustness of the proposed method than the conventional method.

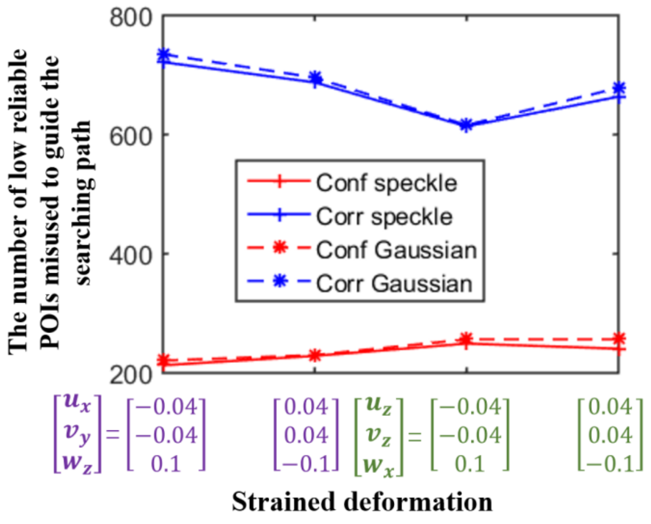


**Fig. S4.** The number of low reliable POIs misused to guide the searching path under various strained deformation in the situation of speckle noise and Gaussian noise.

5. **The proposed confidence weighted strain calculation method vs the conventional strain calculation method**

In fact, the *Conf* can also be used to weight each POI in calculating the strains by the Savitzky-Golay filter. Since the high *Conf* indicates the high reliability of the measurement data of that POI, it is reasonable to assign more weight on the high reliable POI. It is noted that the POI with the *Conf* smaller than the threshold 0.3 is discarded in the strain calculation. Nevertheless, in the conventional method, the strains are directly calculated with the implicit that weights of all the POIs are the same. Fig. S5 shows the strain accuracy comparison between the proposed confidence weighted strain calculation method and the conventional method under various strained deformations in the situation of Gaussian noise and speckle noise. It can be seen that the strains derived from the proposed method are more accurate than those from the conventional method.

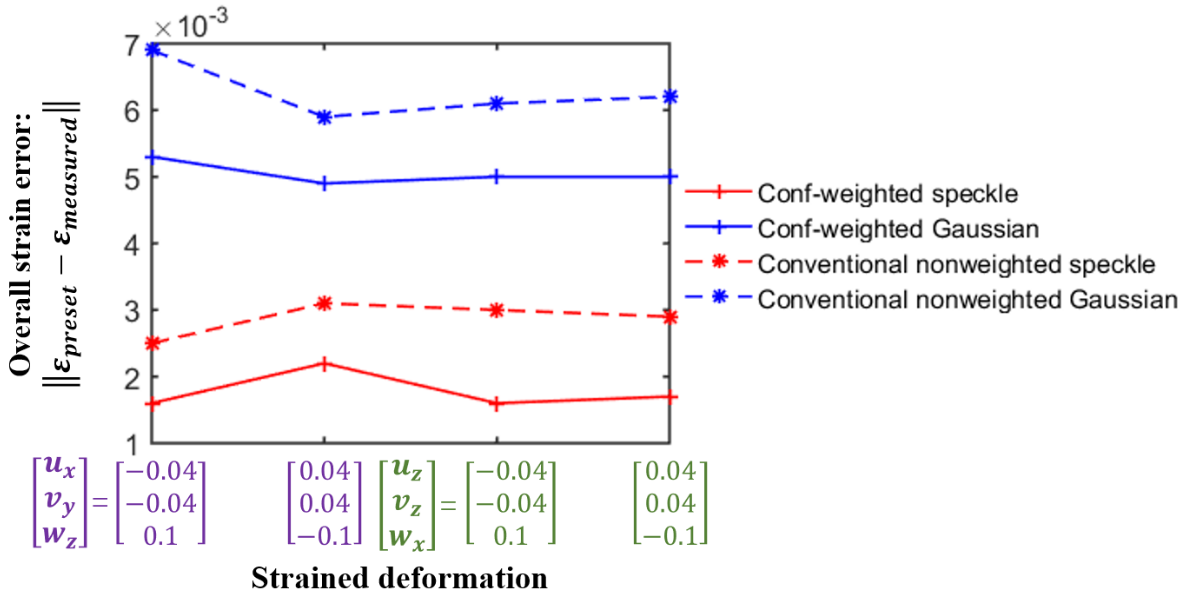


**Fig. S5.** The overall strain errors $\|\varepsilon_{preset} - \varepsilon_{measured}\|$ of the proposed confidence-weighted method and the conventional non-weighted method under various strained deformation in the situation of speckle noise and Gaussian noise.
